# HtsRC-mediated accumulation of F-actin regulates ring canal size during Drosophila melanogaster oogenesis

**DOI:** 10.1101/2020.04.24.060186

**Authors:** Julianne A. Gerdes, Katelynn M. Mannix, Andrew M. Hudson, Lynn Cooley

**Affiliations:** Department of Genetics, Yale University School of Medicine, New Haven, CT; Department of Cell Biology, Yale University School of Medicine, New Haven CT; Department of Molecular, Cellular & Developmental Biology, Yale University, New Haven, CT

## Abstract

Ring canals in the female germline of *Drosophila melanogaster* are supported by a robust filamentous actin (F-actin) cytoskeleton, setting them apart from ring canals in other species and tissues. Previous work has identified components required for the expansion of the ring canal actin cytoskeleton but has not identified the proteins responsible for F-actin recruitment or accumulation. Using a combination of CRISPR-Cas9 mediated mutagenesis and UAS-Gal4 overexpression, we show that HtsRC, a component specific to female germline ring canals, is both necessary and sufficient to drive F-actin accumulation. Absence of HtsRC in the germline resulted in ring canals lacking inner rim F-actin, while overexpression of HtsRC led to larger ring canals. HtsRC functions in combination with Filamin to recruit F-actin to ring-canal-like ectopic actin structures in somatic follicle cells. Finally, we present findings which indicate that HtsRC expression and robust female germline ring canal expansion are important for high fecundity in fruit flies but dispensable for their fertility, a result which is consistent with our understanding of HtsRC as a newly evolved gene specific to female germline ring canals.

## 1 Introduction

Across the animal kingdom, gametogenesis occurs in a syncytium, or a group of interconnected cells (Haglund *et al*. 2011). Germline syncytia are formed through a modified cytokinesis pathway wherein abscission does not occur and daughter cells remain connected by a cytoplasmic intercellular bridge (Haglund *et al*. 2011; Lu *et al*. 2017). In both male and female germlines, intercellular bridges retain some components of the contractile ring, but also recruit additional cytoskeletal proteins to stabilize or grow nascent ring canals (Robinson and Cooley 1996; Greenbaum *et al*. 2007; Haglund *et al*. 2011). However, the process by which intercellular bridges form and recruit components is poorly understood in most species.

Intercellular bridge formation and function are best characterized in the Drosophila ovary where the bridges are called ring canals. In *Drosophila melanogaster*, oogenesis occurs in egg chambers, the functional units of the ovary. Each egg chamber contains a syncytium of 16 germline cells: 15 nurse cells and one transcriptionally quiescent oocyte, formed through four rounds of mitosis via incomplete cytokinesis (Hinnant *et al*. 2020). Following the fourth mitotic division, ring canals increase in diameter, expanding from 0.5-1 *μ*m to a final size of 10 *μ*M or larger, a 20-fold increase (Tilney *et al*. 1996). Ring canals in egg chambers are necessary to support oocyte growth by allowing the flow of cytoplasm (Robinson *et al*. 1994) as well as the transfer of organelles, including mitochondria (Cox and Spradling 2003), from nurse cells to the oocyte. Compared to ring canals in Drosophila males and in females of other species, which range from 1 to 4 μm in diameter, female germline ring canals in Drosophila are the largest ring canals known to date (Haglund *et al*. 2011). Although these ring canals are among the best characterized, the evolutionary and mechanistic reasons for their large size remain a mystery.

The large size of *Drosophila* female germline ring canals depends on the recruitment and expansion of a robust ring canal F-actin cytoskeleton (Tilney *et al*. 1996). After completion of the mitotic divisions, maturing ring canals recruit non-contractile F-actin, HtsRC, a product of the *hu li tai shao (hts)* gene (Yue and Spradling 1992), and Filamin encoded by the *cheerio* gene (Robinson *et al*. 1997). F-actin recruitment requires both HtsRC and Filamin, as the absence of either protein results in F-actin-poor ring canals. The expansion phase is dependent on the activities of the Arp2/3 complex. The Arp2/3 complex acts on existing filaments to promote branching and an increase in the total length of actin filaments at the ring canal (Hudson and Cooley 2002; Zallen *et al*. 2002). Despite playing a role in ring canal expansion, neither the Arp2/3 complex nor Diaphanous is responsible for the initial recruitment of F-actin to ring canals, as mutants and knockdowns of either protein still retain ring canal F-actin. Although Filamin and the Arp2/3 complex are well-characterized regulators of actin polymerization and organization, the functional role of HtsRC and its interactions with existing machinery remains poorly understood.

HtsRC is produced from one class of alternatively spliced mRNAs from the *hts* gene, referred to collectively as *ovhts;* the *hts* gene also produces other splice variants including the conserved F-actin- and Spectrin-binding protein, Adducin (Figure 1A) (Ding *et al*. 1993; Whittaker *et al*. 1999; Petrella *et al*. 2007). Mutations in the *hts* locus were initially isolated from a screen for female sterile mutants; *hts* mutant egg chambers have too few germline cells and ring canals lacking F-actin (Yue and Spradling 1992). The aberrant egg chamber architecture has been attributed to the lack of Adducin, which is required to assemble fusomes needed during early development (Lin *et al*. 1994). An understanding of the contribution of HtsRC to oogenesis has lagged because of its complicated biogenesis and the lack of mutations that affect only the expression of HtsRC.

**Figure 1.**
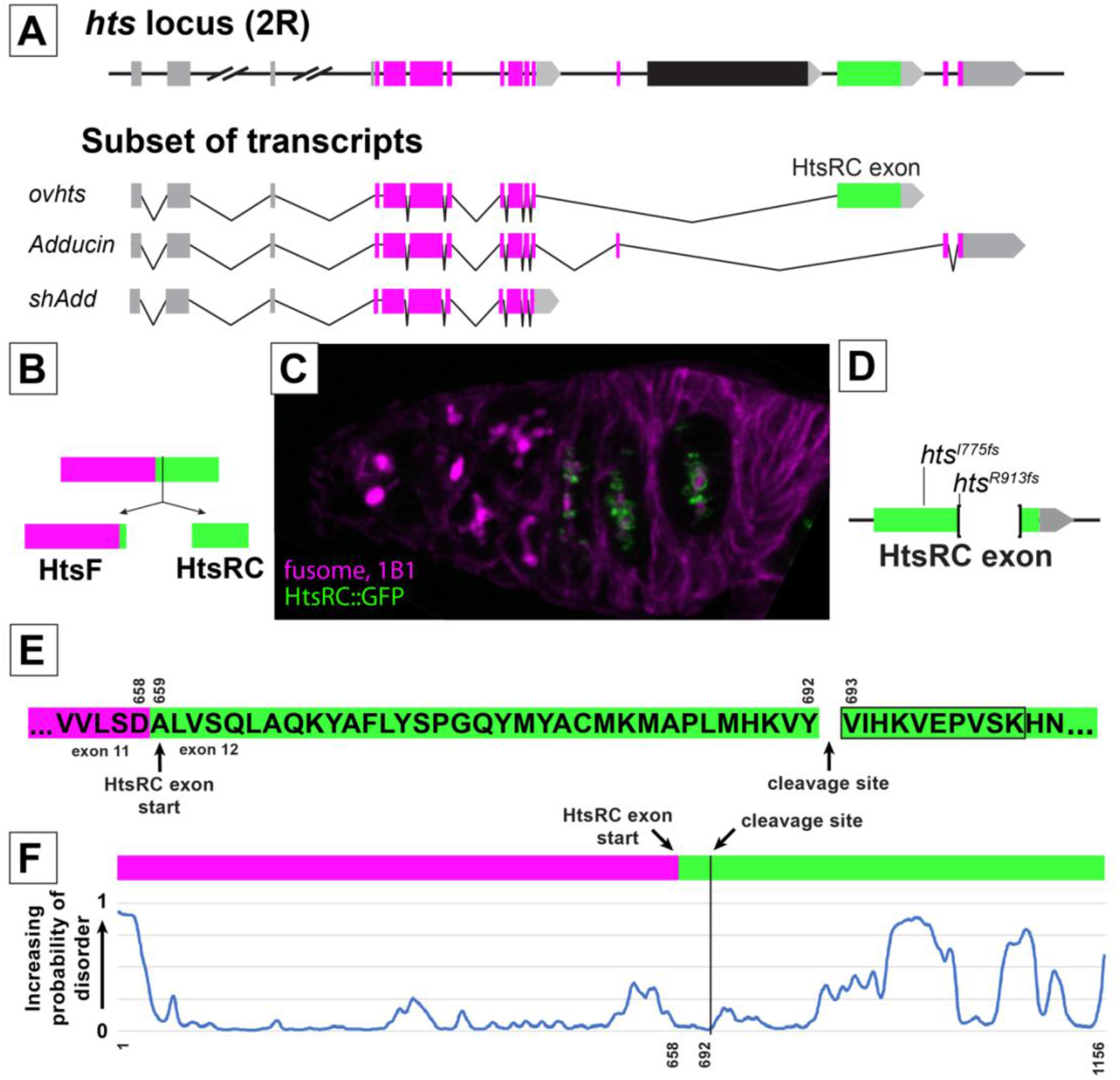
HtsRC is produced from the HtsRC exon after cleavage of the Ovhts polyprotein. (A) Diagram of the *hts* locus, showing a subset of spliced exons (top) and the structure of three transcripts containing alternatively spliced terminal exons. Untranslated regions and exons are color coded as follows: 5’ UTRs (grey boxes), 3’ UTRs (grey arrows), conserved Adducin exons (magenta boxes), and the HtsRC exon (green box). The black exon corresponds to a testis specific splice variant not shown. Three *hts* isoforms with unique terminal exon groups can be detected in the ovary: *ovhts, Adducin*, and *shAdd*. Distances are to scale, except where indicated by hash marks. (B) *ovhts* mRNA produces the Ovhts polyprotein, which is cleaved into HtsF and HtsRC. Exons are color coded as in A. (C) Ovhts polyprotein expressed from the pCOH::ovhts::GFP transgene. Ovhts tagged at the C-terminus with GFP produces HtsRC::GFP and untagged HtsF following cleavage. HtsF is labeled with a monoclonal antibody (1B1) which also labels Add and shAdd in the fusome. (D) Summary of mutations in the HtsRC exon induced by CRISPR-Cas9 mediated NHEJ. *htsI775fs* is a 2 bp deletion resulting in a frameshift and 62 novel amino acid residues followed by a stop codon. *htsR913fs* is a 559 bp deletion followed by a frameshift resulting in 10 novel residues and a stop codon. (E) Amino acid sequence of the Ovhts polyprotein, showing the start of the HtsRC exon and the predicted cleavage site. Mass spectrometry experiments identified fragments of HtsRC with semi-tryptic cleavage sites starting at residue 693 (residue 35 of the HtsRC exon). Black box indicates the identified semi-tryptic peptide. (F) Prediction of intrinsically disordered regions within the Ovhts polyprotein using the SPOT-Disorder-Single application (Hanson *et al*. 2018). Magenta and green bars indicate the HtsF and HtsRC-exon encoded regions respectively. Vertical black line indicates the polyprotein cleavage site described in (E).

The *ovhts* mRNA (Figure 1A) contains a unique terminal exon not present in the *adducin* gene outside of flies (Figure 1A, green exon). In *Drosophila melanogaster*, mRNA containing this exon has only been detected in the ovary, and antibody staining reveals that the protein product of this exon is germline-specific. Previous work from our lab has shown that the Ovhts polyprotein, which is produced from this mRNA, undergoes post-translational cleavage to produce HtsF, a truncated Adducin protein that localizes to fusomes (magenta, Figure 1B-C), and HtsRC, which localizes to ring canals (green, Figure 1B-C) (Petrella *et al*. 2007). In addition to ring canals, HtsRC is also found on cytoplasmic actin bundles at stage 10B and on the oocyte cortex in later stages (Huelsmann *et al*. 2013; Pokrywka *et al*. 2014). The level of HtsRC in egg chambers is under the control of a Cullin3 RING Ubiquitin ligase (CRL3) in which Kelch is the substrate adaptor that binds HtsRC, leading to poly-ubiquitylation of HtsRC and its destruction by the proteasome (Hudson *et al*. 2019). Mutations in the *kelch* gene lead to increased levels of HtsRC and ring canal F-actin, while mutations in the exon encoding HtsRC cause loss of ring canal F-actin, consistent with HtsRC being the target of CRL3_kelch_.

In this work, we investigate how HtsRC controls F-actin levels at the ring canal by testing the impact of loss of function and overexpression of HtsRC. We found that HtsRC is necessary for F-actin accumulation at ring canals and sufficient to drive the aggregation of F-actin when expressed ectopically in somatic follicle cells. HtsRC-dependent F-actin accumulation at ring canals is important for normal ring canal expansion as well as stability, and the F-actin cytoskeleton is essential for robust, high-level fecundity but not fertility. The activity of HtsRC in F-actin accumulation depends on Filamin, but is independent of the Arp2/3 complex. By studying HtsRC-specific mutants, we found evidence for the importance of large stable ring canals in maintaining the high fecundity of *Drosophila melanogaster* females.

## 2 Results

### 2.1 Ovhts polyprotein is cleaved within the HtsRC exon-encoded peptide

We previously identified HtsRC in two independent mass spectrometry approaches designed to identify ring canal-associated proteins (Hudson *et al*. 2019; Mannix *et al*. 2019). Based on our previous work showing that HtsRC is cleaved from the Ovhts polyprotein (Petrella *et al*. 2007), we searched our MS/MS data for peptides resulting from semi-tryptic cleavage, reasoning that an N-terminal peptide beginning at the Ovhts cleavage site could be identified in this way. A search of MS/MS data from proteins that co-purify with the Kelch KREP domain (Hudson *et al*. 2019) identified three peptide spectra with the sequence VIHKVEPVSK (peptide false discovery rate threshold: 0.1%; Figure 1E, black box), consistent with cleavage prior to V693 in Ovhts. The same peptide was identified in a search for semi-tryptic peptides using a dataset from HtsRC::APEX proximity labeling experiments (Mannix *et al*. 2019). We therefore conclude that the Ovhts is cleaved following Y692.

### 2.2 HtsRC is required for the accumulation of F-actin to RCs, but not to other actin-rich structures

Functional characterization of HtsRC in oocyte development was complicated by the important role HtsF and other *hts* isoforms (Figure 1A) play in mitotic divisions and in oocyte specification (Yue and Spradling 1992). Previously isolated mutations in *hts* affected exons common to all *hts* isoforms (Figure 1A, magenta exons), and resulted in female sterility and loss of the fusome in females (Lin *et al*. 1994; Wilson 1999; Petrella *et al*. 2007). To better characterize the functions specific to HtsRC, we used CRISPR-Cas9 gene editing to generate several independent frameshift mutations in the HtsRC-encoding exon, which we initially described in Hudson et al (2019) (Figure 1D). We have designated mutations in the HtsRC exon “*htsRC*-specific mutations” to differentiate them from previously isolated *hts* alleles. Flies homozygous for any of our *htsRC*-specific mutations lack detectable HtsRC protein (Figure 2B” and D”), but have intact fusomes, indicating that Ovhts is processed and cleaved normally and only HtsRC is impacted by the mutations (Hudson *et al*. 2019). F-actin is present in several subcellular locations within the germline cells of egg chambers: ring canals; cell cortexes, including enrichment at microvilli-like structures surrounding ring canals (Loyer *et al*. 2015); and robust cytoplasmic bundles that form specifically during stage 10B (Huelsmann *et al*. 2013; Spracklen *et al*. 2014). Ring canals have an inner rim of highly crosslinked mixed-polarity bundles of F-actin and an outer rim of electron dense plasma membrane (Tilney *et al*. 1996). In *htsRC*-specific mutants, the ring canal inner rim F-actin was completely absent in all stages of egg chamber development (Figure 2A-D), while the ring canal-associated cortical F-actin remained intact (Figure 2D, D’). Filamin, which is present with F-actin and HtsRC in the inner rim (Figure 2A and C), is also associated with the outer rim. Previous data showed that Filamin localization to the outer rim of ring canals is independent of *hts* (Robinson *et al*. 1997; Sokol and Cooley 1999). Similarly, in *htsRC* mutant egg chambers, Filamin was present at outer rims of ring canals lacking inner rim F-actin (Figure 2B-B”, D-D”).

**Figure 2.**
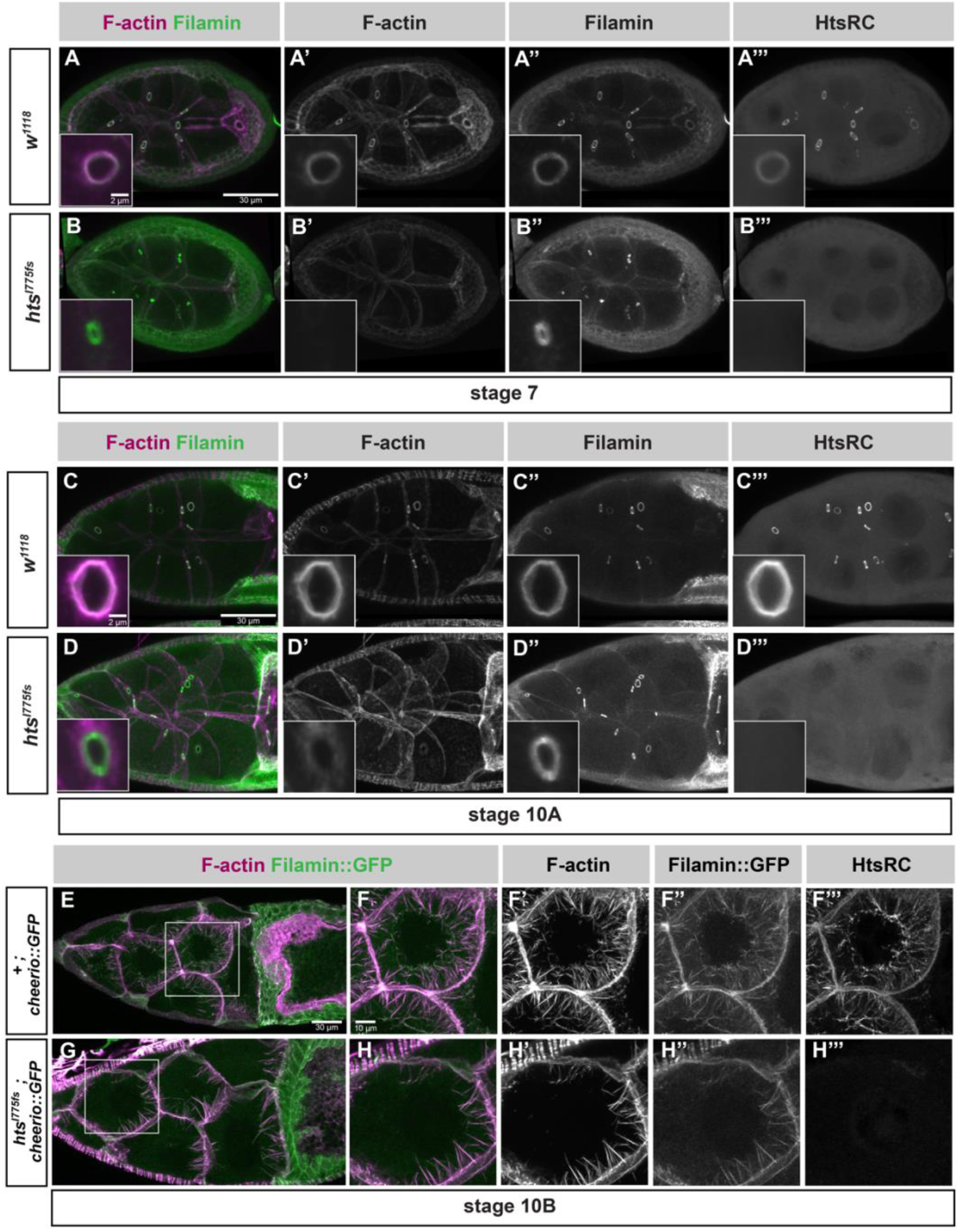
htsRC-specific mutants lose F-actin at ring canals but not at cytoplasmic actin bundles. Maximum intensity projections of wild-type (*w_1118_*) and *htsRC*-specific mutant (*hts_I775_*) egg chambers at (A-B) stage 7, (C-D) stage 10A, and (E-H) stage 10B. (A-D) Egg chambers are stained with TRITC-Phalloidin and labeled with Filamin and HtsRC antibodies. The left column shows a merge of F-actin (magenta) and Filamin (green) channels, where white indicates overlap between channels. Insets contain a representative nurse cell ring canal from the same genotype and stage, although not the same egg chamber. Scale bars are 30 microns, inset scale bars 2 microns. (E-H) Egg chambers from wild-type and *htsRC*-specific mutant flies expressing Filamin::GFP. Size differences between these egg chambers represent the normal variation among stage 10B egg chambers. (E,G) Egg chambers are labeled with TRITC-Phalloidin (magenta). Scale bar is 30 microns. (F,H) Nurse cells marked by white boxes in (E) and (G) are magnified to show labeling of the cytoplasmic actin bundles. Scale bar 10 microns.

Intriguingly, the impact of *htsRC*-specific mutations on F-actin was less severe in the four ring canals connecting the nurse cells directly to the oocyte. In stages 7-10, oocyte ring canals contained some F-actin at the inner rim, which did not occur at nurse cell ring canals (Figure S1). This may be a reflection of differences in the cortical cytoskeleton in the oocyte compared to the nurse cells. Similar robustness at the oocyte cortex has been observed in other studies of ring canal dynamics (Hudson and Cooley 2002; Loyer *et al*. 2015).

HtsRC is also present on cytoplasmic F-actin bundles that form in stage 10B nurse cells (Huelsmann *et al*. 2013); however, it was not known whether HtsRC is required for bundle formation. The F-actin bundles are monopolar, with plus ends at the plasma membrane and minus ends that reach the nuclear envelope as the bundles lengthen. The bundles contain at least three F-actin crosslinking proteins - Fascin encoded by the *singed* gene (Cant *et al*. 1994; Cant and Cooley 1996), Villin encoded by the *quail* gene (Mahajan-Miklos and Cooley 1994), and Filamin encoded by the *cheerio* gene - along the length of the bundles, and HtsRC protein concentrated at the plus ends (Huelsmann *et al*. 2013). Flies with strong mutations in *singed* or *quail* are sterile due to a failure to form F-actin bundles, resulting in ring canals plugged by the large nurse cell nuclei during nurse cell dumping in stage 10B and 11 egg chambers (Cant *et al*. 1994; Mahajan-Miklos and Cooley 1994). In contrast, stage 10B egg chambers in *htsRC*-specific mutants produced cytoplasmic F-actin bundles containing Filamin::GFP (Fig 2E-H) and nurse cell nuclei did not interfere with RCs. This result suggests that HtsRC is not essential for cytoplasmic F-actin bundle formation late in oogenesis.

We also tested whether HtsRC localization at cytoplasmic F-actin bundles depends on Filamin. In three independent *cheerio* mutants, HtsRC protein localized to the bundles as in wild type (Figure S3). This was consistent with observations of protein localization on cytoplasmic bundles that did not reach the nucleus: HtsRC was present on bundles while Filamin was found at the nuclear envelope and these proteins only colocalized when cables successfully reached the nucleus (Huelsmann *et al*. 2013). This is different from ring canals where HtsRC localization is dependent on Filamin (Robinson *et al*. 1997), perhaps because the presence of other F-actin cross-linking proteins can compensate for the absence of Filamin in cytoplasmic bundles.

### 2.3 Ring canals lacking F-actin are small and unstable

Ring canal size is known to depend on the recruitment and elongation of F-actin. Previous work characterizing the role of the Arp2/3 complex in ring canal expansion showed ring canal diameter decreased by 30 to 50% in *ArpC1_Q25st_* and *Arp3_EP(3)3640_* mutant clones, where truncation of the ArpC1 subunit or absence of the Arp3 subunit rendered the Arp2/3 complex inactive (Hudson and Cooley 2002). Ring canals in *ArpC1_Q25st_* and *Arp3_EP(3)3640_* mutant cells still contained F-actin and the first point at which they had a significant reduction in size was stage 5, the point at which rapid ring canal expansion begins (Tilney *et al*. 1996; Hudson and Cooley 2002). As our *htsRC*-specific mutants failed to recruit F-actin at any stage, we expected to see a similar decrease in ring canal size in our *htsRC*-specific mutants.

Filamin persists on ring canals in HtsRC mutants, allowing us to measure ring canal diameter. In *htsRC*-specific mutants, ring canal diameter was smaller compared to wild type, but only for RCs connecting two nurse cells (Figure 3A-D). The difference in nurse cell ring canal diameter was easily observable starting at stages 4 and 5, but was statistically significant even at stage 2 (Figure 3C). At stage 2, the *htsRC-specific* mutants had nurse cell ring canal diameters that were 18% smaller than in wild type on average (1.4 micron vs. 1.7 microns); after stage 5, the diameter of these ring canals was reduced by 30 to 40% compared to wild type. By stage 10B, the average nurse cell ring canal in *htsRC*-specific mutants was 5.2 microns in diameter, compared to 9.0 microns in wild-type controls. Unlike nurse cell to nurse cell ring canals, the four ring canals connecting nurse cells to the oocyte did not show a significant change in diameter (Figure 3D). The difference we measured in stage 10A (Figure 3D) may have been due to the wide range of ring canal sizes during stage 10. The heterogeneity between nurse cell and oocyte ring canal phenotypes was consistent with the results reported for *ArpC1_Q25sd_* (Hudson and Cooley 2002), and may be related to the residual F-actin we saw at oocyte ring canals (Figure S1). Although there was not a significant decrease in size for oocyte ring canals, they did exhibit a statistically significant change in variance, consistent with the presence of both extremely large and extremely small ring canals in that data set (Figure 3D). Our results for *htsRC* mutants, combined with previous results for the Arp2/3 complex, strongly suggest that the ring canal F-actin cytoskeleton is required for ring canal expansion. HtsRC functions to recruit F-actin and Arp2/3 promotes F-actin nucleation and elongation.

**Figure 3.**
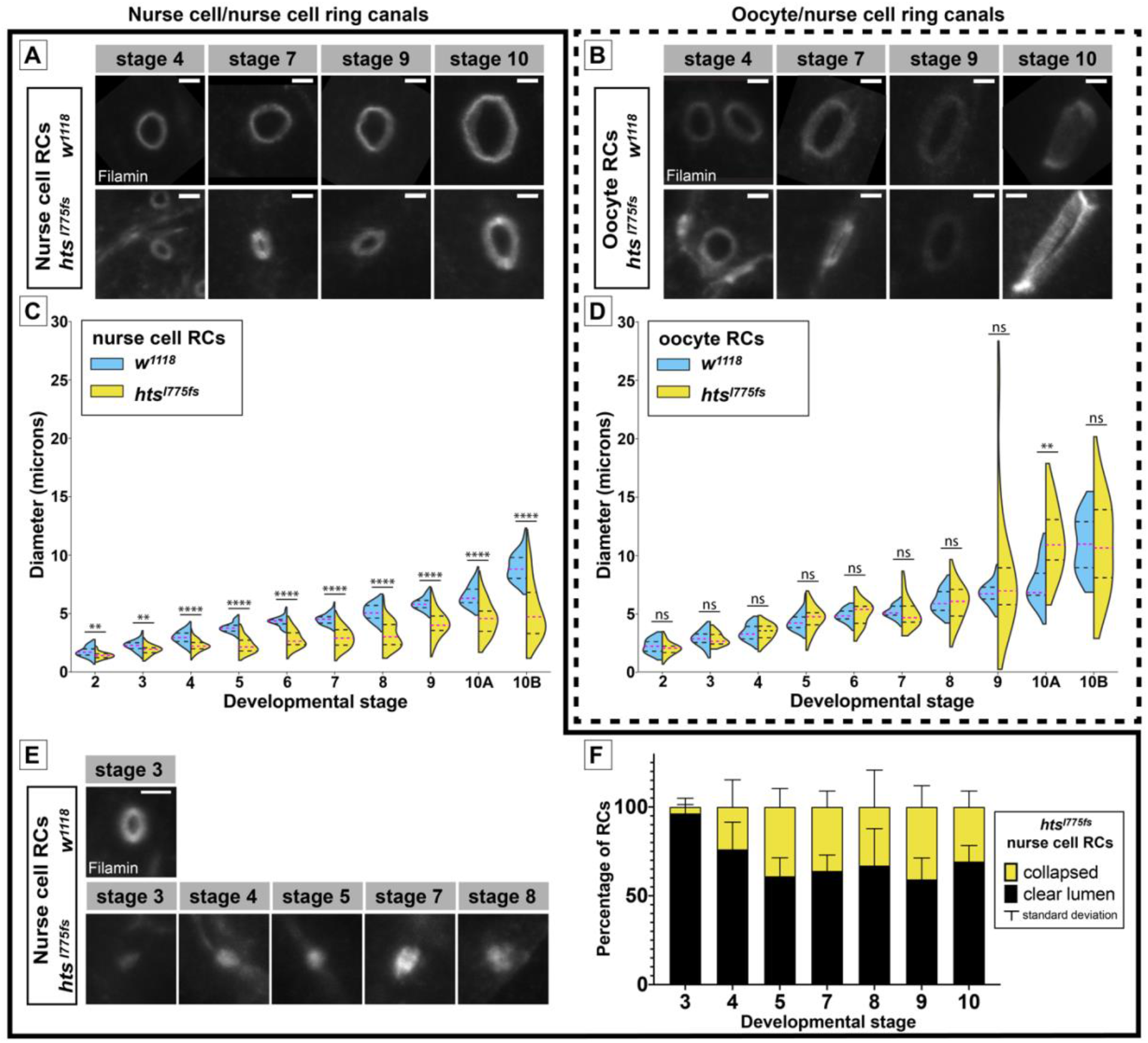
Nurse cell ring canals are smaller and less stable in HtsRC mutants. Representative ring canals and quantification of ring canal diameter for ring canals connecting pairs of nurse cells (A,C,E) or connecting nurse cells to the oocyte (B,D). (A-B) Representative ring canals labeled with Filamin antibody from *w_1118_* (top) and *hts_I775_* (bottom) ovaries. (C-D) Violin plots showing ring canal diameter across stages 2-10B of oogenesis in *w_1118_* (blue) vs *hts_I775_* (yellow). Two or more egg chambers were sampled at each stage (22-77 nurse cell ring canals, 8-28 oocyte ring canals). Significance was determined by Welch’s t-test. Significance thresholds: **, p < 0.01; ****, p < 0.0001; ns, not significant. (E) Representative examples of collapsed nurse cell ring canals in *htsRC*-specific mutants (bottom) compared with a wildtype *w1118* ring canal (top) with a clear lumen. (F) Number of collapsed nurse cell ring canals (no visible lumen; yellow) compared to nurse cell ring canals with a clear open lumen (black). Each egg chamber contains 11 nurse cell ring canals. n=3 egg chambers per stage. Scale bars are two microns.

In addition to the effects on ring canal diameter, a subset of ring canals in *htsRC*-specific mutants had no visible lumen and appeared collapsed into foci that formed a single peak on a line scan of pixel intensity (Figure 3E). Collapsed ring canals were present starting at stage 3-4; some stage 3 egg chambers did not contain a single collapsed ring canal, while every stage 4 or older egg chamber contained at least one. This suggests that ring canals formed but grew slowly and sometimes collapsed, rather than failing to expand at all. No collapsed oocyte ring canals were observed, consistent with the more stable nature of oocyte ring canals.

### 2.4 Defective cytoplasm transfer causes fecundity defects in *htsRC-specific* mutants

Previously studied mutations in *hts*, as well as mutations in other ring canal components, namely *kelch* and *cheerio* (Filamin), resulted in completely sterile female flies. In *kelch* and *cheerio* ovaries, egg chambers progressed to stage 11 when the final rapid transfer of nurse cell cytoplasm to the oocyte was hampered by mutant ring canals, resulting in “dumpless” egg chambers containing residual nurse cells that produced small, infertile eggs (Xue and Cooley 1993; Robinson *et al*. 1994, 1997). The phenotype of *hts* null mutant egg chambers was more severe with arrested development and degeneration of early egg chambers (Yue and Spradling 1992).

Surprisingly, *htsRC*-specific mutants were fertile; however, egg chamber development was not normal and hatching of eggs laid by mutant females was markedly reduced. Approximately one quarter of oocytes were the same size and shape as in wild type, while the remaining three quarters of egg chambers showed a variably penetrant dumpless phenotype which we quantified by measuring egg length for two independent *htsRC*-specific mutants (Figure 4A). Eggs laid by *htsRC* mutant females ranged in size from full-sized (>0.5 mm) to less than half the length of wild-type eggs (<0.25 mm; Figure 4A-B). Both egg laying and hatching rates were significantly decreased in *htsRC*-specific mutants (4C-E). Although most wild-type eggs hatched (86%), eggs in two independent *htsRC*-specific mutants hatched less than 30% of the time, and small eggs (produced from dumpless egg chambers) never hatched (Figure 4E). These results were consistent for both *hts_I775fs_* and *hts_R913fs_* alleles and for homozygotes and flies hemizygous for mutant alleles over a deficiency, leading to the conclusion that HtsRC is required for normal levels of fecundity.

**Figure 4.**
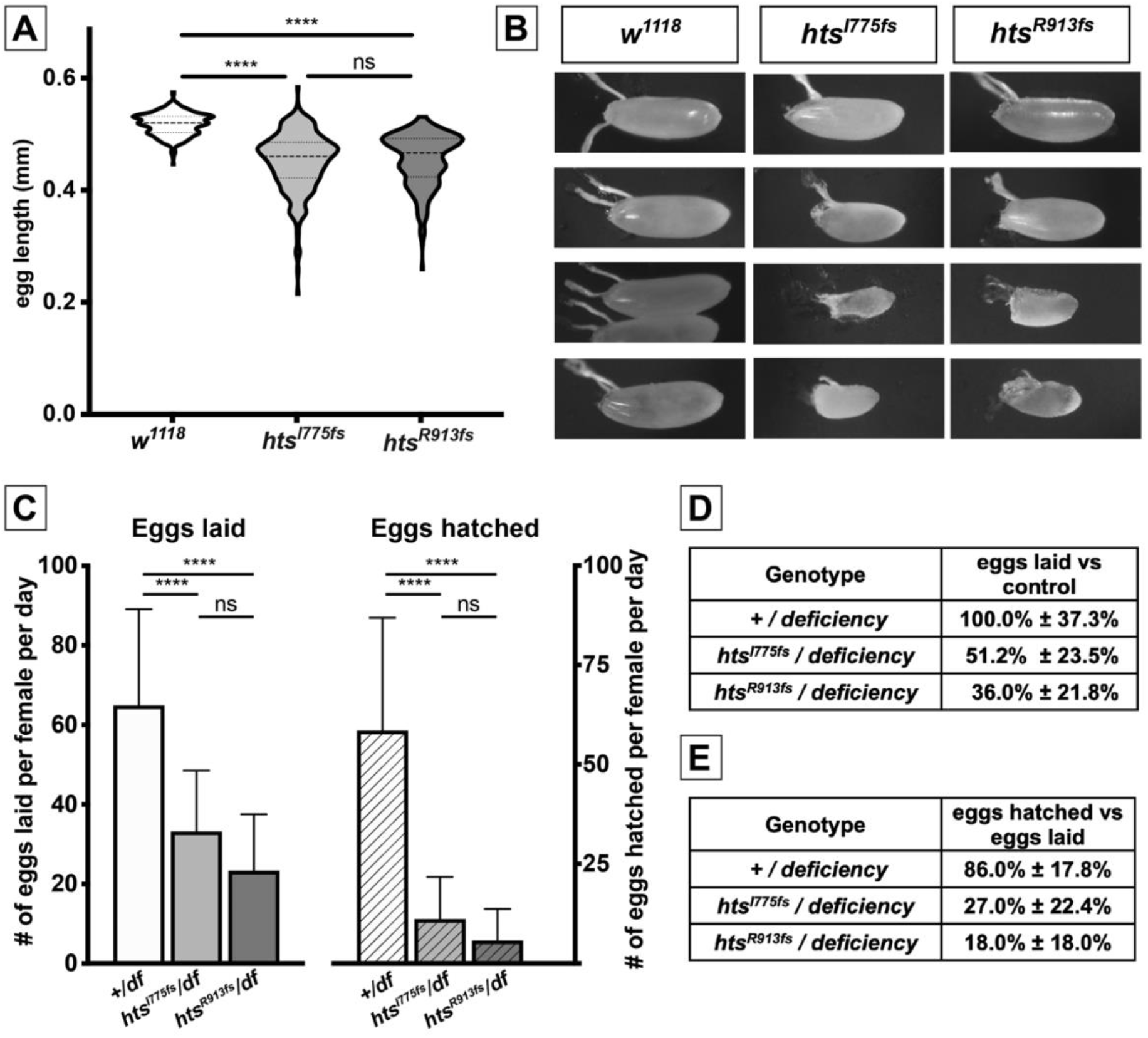
Loss of HtsRC protein impacts egg length and fecundity. (A) Violin plots showing distribution of egg length for two unique *htsRC*-specific mutants or *w_1118_*. n>200 per genotype. Significance was determined using Tukey’s multiple comparison test. (B) Representative images showing the range of sizes among laid eggs. (C) Graph indicating number of eggs laid per day per female (solid bars) compared with the number of eggs hatched per female (patterned bars) for flies carrying either a wild type allele or one of two independent *htsRC*-specific mutations over the *Df(2R)BSC26* deficiency. Error bars indicate standard deviation. n>11 per genotype. Significance was determined using Tukey’s multiple comparison test. (D) Egg laying rates with standard deviation as a percentage of egg laying in the wild type control. (E) Egg hatching as a percentage of the number of eggs laid for each genotype, +/- standard deviation. Significance thresholds: ****, p < 0.0001; ns, not significant.

### 2.5 Overexpression of HtsRC in the germline drives increased RC diameter

As loss of HtsRC resulted in smaller ring canals, we sought to further investigate the role of HtsRC in the regulation of ring canal expansion by examining the effects of HtsRC overexpression. To achieve overexpression, *mat–GAL4* was used to drive *ovhts::GFP* in stage 2 and older egg chambers. Ovhts polyprotein produced by this transgene undergoes complete cleavage to produce functional HtsRC protein that is detectable with HtsRC antibody or by GFP fluorescence (Petrella *et al*. 2007). In control egg chambers (*w_1118_*), ring canal expansion occurred gradually until reaching peak size at Stage 10B (Figure 5A, A’, C, C’, E, E’). In contrast, overexpression of HtsRC resulted in larger ring canals at all stages of oogenesis examined (Figure 5B, B’, D, D’, F, F’). The increase in ring canal diameter was observed in both nurse cell (Figure 5G-H) and oocyte (Figure 5I-J) ring canals. Quantification of ring canal diameter in control and HtsRC-overexpression backgrounds revealed that the differences in ring canal sizes were statistically significant throughout oogenesis and became more apparent in late-stage egg chambers (Figure 5K). As ring canal growth is driven by F-actin polymerization, these results suggest that HtsRC contributes to ring canal expansion, presumably by working to recruit and/or regulate the ring canal F-actin cytoskeleton.

**Figure 5.**
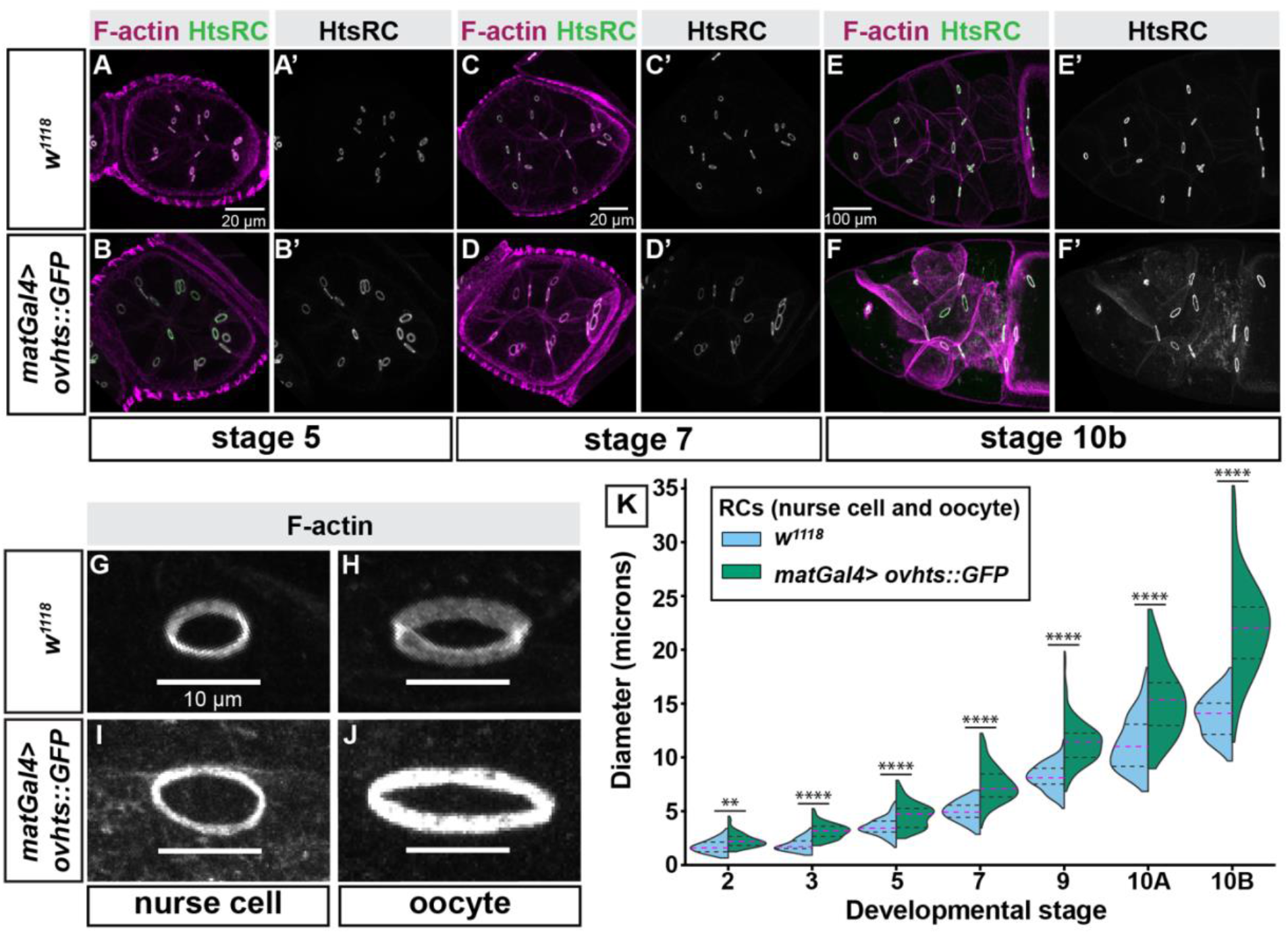
Overexpression of *ovhts* results in larger ring canals. (A-F) Fluorescence micrographs showing F-actin and HtsRC antibody stain in egg chambers throughout oogenesis show that *maternal alpha tubulin–GAL4* (*matGal4*)–driven *UASH-ovhts::GFP* (with *htsN43’UTR)* leads to the formation of large ring canals, compared to *w_1118_* control. (G-J) Higher-resolution micrograph of nurse cell and oocyte ring canals of a stage 10 egg chamber stained with F-actin to show the increased diameter of the matGal4>*ovhts::GFP* ring canals compared to the *_w1118_* control. (K) Quantification of ring canal diameter throughout development, visualized by violin plots. Magenta dotted line indicates median ring canal diameter, and black dotted lines represent the upper and lower quartiles. Welch’s t-test was performed between *_w1118_* control and *matGal4>ovhts::GFP* ring canal diameters at each stage; Significance thresholds: **, p < 0.01; ****, p < 0.0001.

### 2.6 HtsRC ectopically expressed in somatic follicle cells drives the formation of F-actin aggregates that mimic the ring canal actin cytoskeleton

Genetic analysis provided strong evidence that HtsRC functions in F-actin regulation. Efforts to investigate the biochemical activity of HtsRC using standard biochemical assays of F-actin polymerization and bundling were unproductive. Numerous attempts at expression and purification of HtsRC were unsuccessful, most likely because of the intrinsically disordered nature of the HtsRC protein. Seeking an alternative approach, we developed an ectopic expression system in which to further test HtsRC function.

We expressed HtsRC ectopically in the somatic epithelial cells of egg chambers, which are polarized with the apical side in contact with the germline (Figure 6A-C) (Horne-Badovinac and Bilder 2005). The cortex of follicles cells is F-actin-rich, allowing it to be visualized using Phalloidin (Fig 6B’, C’). Follicle cells develop in syncytia of variable size that remain connected by ring canals about 1 *μ*m in diameter located near the apical end (McLean and Cooley 2013). HtsRC is not normally expressed in these cells (Figure 6B”, C”) and their ring canals do not contain Filamin or Kelch, although they do contain Pavarotti, which has been found on all known RCs (Lu *et al*. 2017).

**Figure 6.**
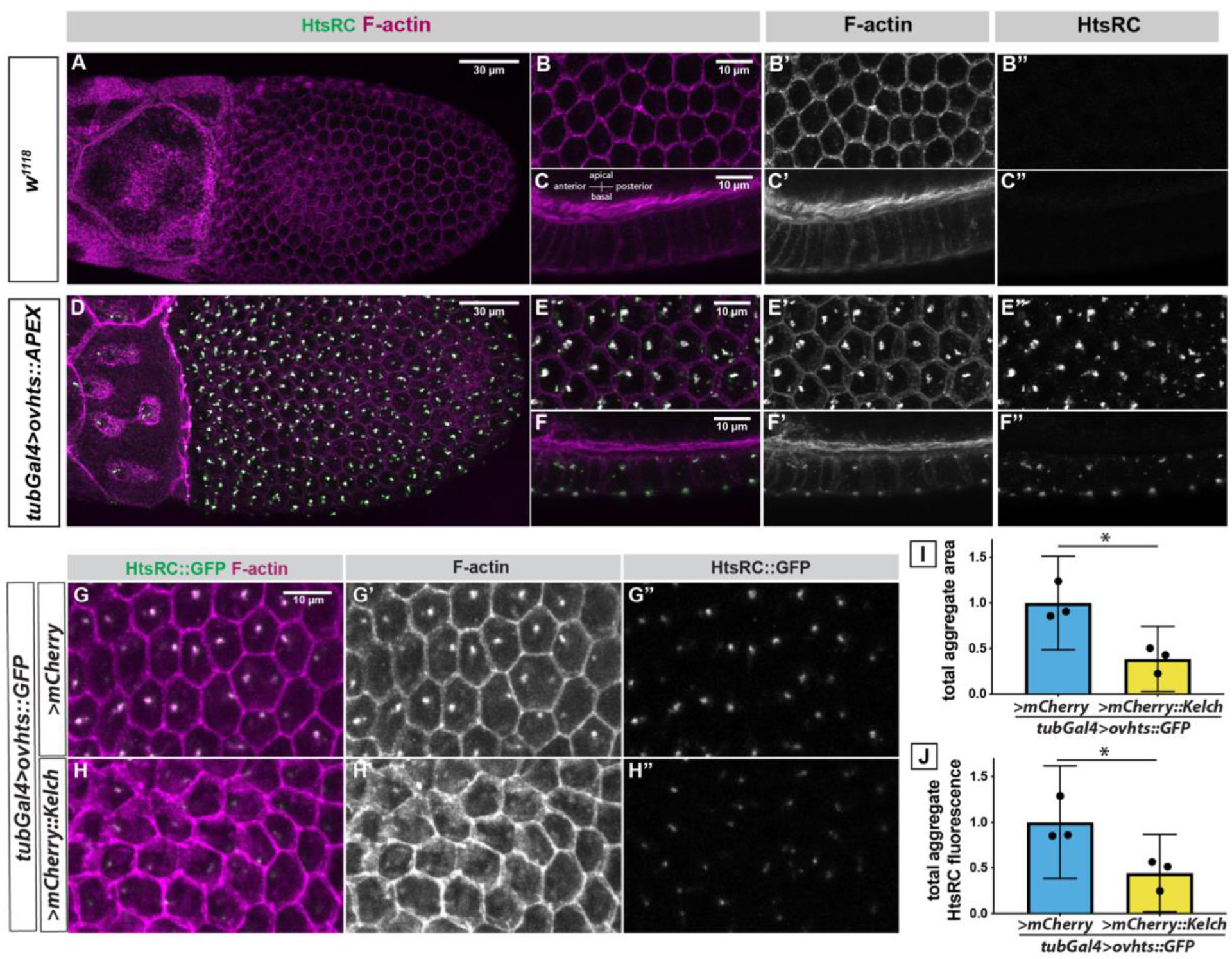
Ectopic HtsRC expression drives the formation of F-actin aggregates. (A-F) Stage 10B egg chambers focusing on the somatic follicle cell layer surrounding the oocyte either from the basal surface (A, B, D, E) or from the side with the basal side at the bottom and the apical side and germline at the top (C, F). (A-C) Wildtype (*_w1118_*) control egg chamber labeled with Phalloidin and HtsRC antibody. All images shown were taken from the same egg chamber. (D-F) Ectopic expression of *UASh-ovhts::APEX::V5* driven with the *tubulin-Gal4 (tubGal4)* driver. All images are taken from the same egg chamber. (G-H) *tubulin-Gal4* driving *UASh-ovhts::GFP* along with either *UASp-mCherry::kelch* or a *UASp-mCherry* control. (I) Total area occupied by foci in a Z-projection the follicle cells, measured by HtsRC::GFP fluorescence. (J) Total HtsRC::GFP fluorescence contained within aggregates in control or *UASp-mCherry::kelch* overexpression. (I-J) Student’s t-tests were used to determine significance. Significance thresholds: *, p < 0.05; **, p < 0.01; ***, p < 0.001; ****, p < 0.0001; ns, not significant.

When *ovhts* cDNA transgenes were expressed in the somatic follicle cells using the *tubulin-Gal4* driver, prominent aggregates containing HtsRC and F-actin were visible (Figure 6D-F). The aggregates always formed near the cortex of these cells, with most of them near basal or apical surfaces (Figure 6F). Smaller aggregates were also present near the lateral cortex and on the basal and apical survaces. Each cell contained at least one basal aggregate, but the number and location of the other aggregates were variable. We tested the ability of several HtsRC-expressing transgenes to cause the formation of aggregates, including transgene constructs containing different untranslated regions and tags, and either full length *ovhts* cDNA or just the HtsRC-encoding exon (not shown). The location and frequency of aggregate formation remained consistent, suggesting their formation was driven by HtsRC, and not by aggregation of GFP or APEX protein tags (Figure S3).

HtsRC did not colocalize with follicle cell ring canals as we initially predicted; however, HtsRC/F-actin aggregates did contain several components of ovarian ring canals, including Kelch, Filamin, and Arp3 suggesting they could be used as a proxy for ring canals to analyze HtsRC function. We have shown that Kelch regulates HtsRC levels by targeting it for destruction by the proteasome in the germline (Hudson *et al*. 2019). To test whether HtsRC was similarly regulated by Kelch in follicle cells, we co-expressed Ovhts::GFP with either mCherry as a control or mCherry::Kelch (Figure 6G-H). In the presence of additional Kelch, the number of aggregates and overall aggregate area were reduced (Figure 6I). F-actin levels were also reduced relative to the control (Figure 6J). These data suggest that HtsRC in ectopic aggregates undergoes Kelch-dependent turnover, as it does in germline ring canals.

To further test the impact of Kelch on HtsRC, we co-expressed an overactive form of Kelch lacking its N-terminal regulatory sequence, Kelch_ΔNTR_ (Figure S4). In this background, HtsRC/F-actin aggregates were almost entirely absent, consistent with our previous results in germline cells (Hudson *et al*. 2019). These results indicate that ectopically expressed HtsRC can be regulated in a manner similar to its regulation in the female germline, making HtsRC-induced follicle cell aggregates an appropriate system for probing HtsRC function.

The ability of HtsRC to cause the assembly of F-actin structures in germline ring canals and somatic follicle cells likely involves other proteins. We tested whether the Arp2/3 complex is needed for HtsRC-induced F-actin in follicle cells. The Arp2/3 complex is required for germline ring canal expansion beginning at stage 5, but not for the initial recruitment of F-actin to ring canals. In germline clones homozygous for *ArpC1_Q25st_* or *Arp3_EP(3)3640_*, null mutations in components of the Arp2/3 complex, HtsRC and F-actin were still present on ring canals (Hudson and Cooley 2002). These data suggested that HtsRC may act upstream or independently of the Arp2/3 complex to recruit or regulate F-actin in the context of a ring canal.

To determine if HtsRC can assemble F-actin in the absence of Arp2/3 complex activity, we examined the effect of loss of Arp2/3 activity in follicle cells using the MARCM system (Lee and Luo 1999). Large clones of *ArpC1_Q29sd_* or *ArpC4S_H1036_* mutant cells expressing the HtsRC::APEX transgene contained F-actin aggregates with the same size and localization pattern as neighboring heterozygous cells (Figure S4). These results suggest that HtsRC generates F-actin rich structures independently of the Arp2/3 complex in both germline and follicle cells.

### 2.7 Filamin is important for HtsRC-mediated F-actin aggregate formation

Filamin is required for recruitment of HtsRC to ring canals (Robinson *et al*. 1997) and more recent work in our lab identified Filamin as a nearest neighbor and likely interaction partner for HtsRC (Mannix *et al*. 2019). However, it is not possible to test the function of HtsRC independently of Filamin at the ring canal, as ring canals do not form or recruit HtsRC in *cheerio* mutants (Robinson *et al*. 1997). By leveraging the HtsRC/F-actin aggregates in somatic follicle cells, we were able to separate HtsRC-dependent recruitment of F-actin from ring canal formation.

To test the role of HtsRC independently of Filamin, we examined the formation of F-actin aggregates in *cheerio* mutant somatic follicle cells overexpressing HtsRC from the *tubulin-ovhts-htsN4_Δ100_* transgene. This *ovhts* construct was previously shown to drive the formation of HtsRC/F-actin structures (Pokrywka *et al*. 2014). We have found the structures formed by this transgene in the somatic follicle cells to be consistent with those generated by other HtsRC-expressing transgenes driven by *tubulin-Gal4* (Figure S3). In control follicle cells, HtsRC-positive F-actin aggregates were readily observed at the basal surface (Figure 7A-A”) as well as at the apical and lateral membranes (Figure 7B-B”). In comparison, F-actin aggregates were largely disrupted in the absence of Filamin, as shown with two independent *cheerio* alleles (Figure 7C-F”). To quantify the phenotype, we measured the intensity of HtsRC antibody labeling and phalloidin staining within each aggregate in a defined region and derived a density value by dividing by the volume of the aggregate; the sum of all aggregated densities was calculated for each region (see Materials and Methods). We observed a decrease in the density of both HtsRC (Figure 7G) and F-actin (Figure 7H) in the absence of Filamin. Furthermore, there was less F-actin per unit of HtsRC in the aggregates (Figure 7I), suggesting the ability of HtsRC to recruit F-actin is diminished in the absence of Filamin.

**Figure 7:**
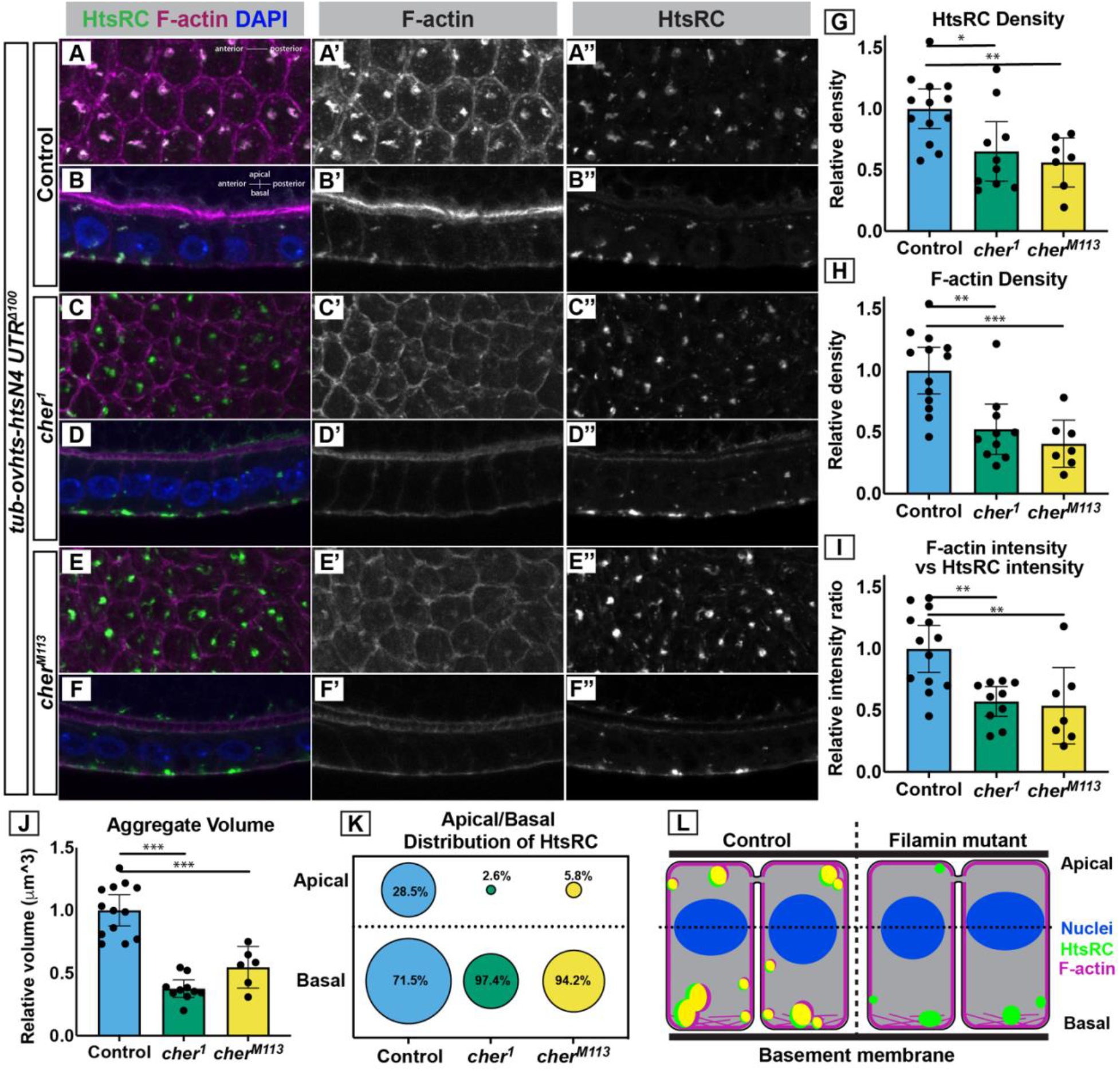
Filamin is important for F-actin recruitment to HtsRC aggregates. Ectopic HtsRC protein was driven from the *tubulin-ovhts-N4_△100_* transgene in somatic follicle cells in either a wildtype background (A-B) or two independent *cheerio* mutant backgrounds (C-F). (A,C,E) Z-projections from representative egg chambers showing the follicle cells from the basal surface. (B,D,F) Z-projections of follicle cells showing the apical/basal distribution of aggregates with the apical side facing the top. Image pairs come from the same egg chamber. (G-H) Density calculated as HtsRC or F-actin density per micron3, relative to the mean of the control. (I) F-actin intensity per unit HtsRC intensity, relative to the mean of the control. (J) Total volume occupied by aggregates normalized to mean aggregate volume in control. (G-J) All results were significant by both ANOVA and pairwise t-tests, with pairwise T-tests displayed on graphs. Significance thresholds: *, p < 0.05; **, p < 0.01; ***, p < 0.001. (K) Distribution of aggregates relative to the center of the nuclei (dotted line in L). Percentages indicate the distribution of HtsRC as a percentage of total HtsRC intensity within that genotype. The areas of the circles are scaled to indicate relative intensity between genotypes and between. (L) Model of HtsRC/F-actin aggregate formation in control (left) compared to *cheerio* mutants (right). HtsRC (green) and F-actin (magenta) colocalize (yellow) in aggregates at both the apical and basal ends of the cells in controls, but HtsRC does not recruit F-actin robustly in mutants and primarily forms aggregates at the basal end.

The total volume of aggregates was also markedly reduced in *cheerio* mutant follicle cells (Figure 7J), with the greatest effect on the apical population of aggregates, which were nearly absent (Figure 7K). Like the oocyte cortex in the germline, the basal cortex of the follicle cells has a greater enrichment of F-actin than the other surfaces in this cell type (Gutzeit 1990), which may contribute to the persistence of basal aggregates in the absence of Filamin.

## 3 Discussion

In this work, we have determined that HtsRC is essential for the recruitment or retention of F-actin at ring canals. In the absence of HtsRC, other F-actin regulators including Filamin and the Arp2/3 complex were insufficient to promote ring canal F-actin accumulation and ring canal expansion. When HtsRC-dependent accumulation of F-actin was disrupted, ring canals did not fully expand. Although some ring canals in *htsRC*-specific mutants were unstable and collapsed, the lumens of most ring canals remained clear. Unlike most other mutations impacting ring canals, the loss of HtsRC did not render females sterile. Instead, cytoplasm transfer to oocytes was compromised and fecundity was reduced. We also found evidence that HtsRC is a potent driver of F-actin recruitment when expressed ectopically, and that Filamin function is linked to HtsRC’s ability to accumulate F-actin. In egg chamber germline cells, Filamin is required for HtsRC localization to ring canals (Robinson *et al*. 1997). In somatic follicle cells, the lack of Filamin impairs the ability of HtsRC to accumulate F-actin in ectopic structures. We conclude that HtsRC is a potent actin regulator and the primary driver of ring canal F-actin accumulation in egg chamber germline cells, and that the recruitment of F-actin is important, but not essential for ring canal stability and fertility.

### 3.1 HtsRC functions with Filamin to promote ring canal F-actin accumulation

In this work, we have shown that HtsRC stimulates the accumulation of F-actin at ring canals and in ectopic structures, and uncovered evidence that helps to place the function of HtsRC in the context of known F-actin regulatory proteins. In *htsRC*-specific mutants, F-actin is not accumulated at ring canals. Based on this phenotype, the function of HtsRC could be to nucleate new actin filaments at the ring canal or to stabilize filaments recruited to ring canals; however, the localization of HtsRC points to a role in stabilizing F-actin. Although HtsRC colocalizes ubiquitously with the mixed-polarity bundles at ring canals, it preferentially binds F-actin filaments near pointed ends in two contexts: the cytoplasmic bundles in nurse cells (Figure 2) (Huelsmann *et al*. 2013; Spracklen *et al*. 2014) and ectopic F-actin filaments formed at the oocyte cortex during later stages of oogenesis (Pokrywka *et al*. 2014). This localization pattern is similar to that of cofilin, which has been shown to preferentially interact with filaments containing ADP-actin at the pointed end of filaments (Carlier *et al*. 1997). At high concentrations, cofilin coats F-actin filaments and stabilizes them (Andrianantoandro and Pollard 2006; Ngo *et al*. 2015); HtsRC could function similarly. Although actin nucleation factors including Arp2/3 and Spire also interact with the pointed ends of filaments, these factors are retained at the end of the filament following nucleation (Mullins *et al*. 1998; Quinlan *et al*. 2005) and HtsRC coats the sides of filaments. Furthermore, the absence of HtsRC specifically at the barbed ends of nurse cell actin bundles (at the plasma membrane) also argues against a role for HtsRC as a nucleation or elongation factor, especially because it lacks domains characteristic of nucleation factors such as formin homology domains or tandem arrays of WH2 domains (Dominguez 2009). In addition, the F-actin aggregates formed by ectopic expression of HtsRC in follicle cells form only at the F-actin-rich cortex, most prominently the basal surfaces, suggesting it requires existing filaments to initiate aggregate formation. HtsRC also requires Filamin to promote robust accumulation, consistent with a possible interaction between HtsRC and existing filaments bound by Filamin.

We propose that HtsRC functions as a scaffold for the ring canal F-actin cytoskeleton. Fluorescence Recovery After Photobleaching (FRAP) experiments showed that GFP-tagged F-actin recovers rapidly with a t1/2 of 65 seconds (Kelso *et al*. 2002) while GFP-tagged HtsRC does not recover in 30 minutes (L. Petrella and L. Cooley, unpublished), suggesting HtsRC could be part of a stable matrix for collecting actin filaments. We also know that the levels of HtsRC are regulated by ubiquitin-mediated proteasome turnover orchestrated by CRL3_Kelch_. Since most of the HtsRC at the ring canal is stable to turnover, CRL3_Kelch_ is likely acting on a subset of HtsRC in order to keep the lumen clear of F-actin as the ring canal grows. We determined that CRL3_Kelch_ also controls HtsRC expressed in follicle cells, suggesting that HtsRC’s potent F-actin accumulation activity is restricted to specific cellular locations where it is resistant to CRL3_Kelch_-mediated turnover. Overall, our results point to recruitment or stabilization of existing filaments as the function of HtsRC.

Our genetic results support a strong functional link between HtsRC and Filamin in ring canals and when HtsRC is expressed ectopically. Our proximity labeling experiments support the conclusion that HtsRC and Filamin are direct binding partners (Mannix *et al*. 2019). In ring canals, Filamin localizes to nascent ring canals after the fourth and final mitotic division of cystocytes independent of HtsRC. The simplest model is that Filamin directly recruits HtsRC, and HtsRC recruits F-actin. How Filamin localizes to new ring canals is not known. Its large size and elongated structure have been shown to mediate many protein interactions in numerous cellular processes (Nakamura *et al*. 2011; Razinia *et al*. 2012) and it may function to link HtsRC and other ring canal components to the membrane. Filamin also has multiple actin binding domains and is known to facilitate the formation of both F-actin bundles and orthogonal F-actin structures (Nakamura *et al*. 2007). Even if HtsRC is the primary recruitment factor at ring canals, the behavior of HtsRC in follicle cells suggests that the presence of Filamin may enhance HtsRC activity in promoting F-actin structures. In follicle cells, HtsRC-dependent actin structures are less robust without Filamin, with smaller aggregates containing less HtsRC that recruit less F-actin per unit of HtsRC. Filamin may function to remodel ring canal F-actin recruited by HtsRC into condensed and ordered structures. Alternatively, HtsRC itself may depend on an interaction with Filamin to stimulate its activity, possibly by changing the structure of HtsRC protein to a more active conformation or by sequestering HtsRC away from CRL3Kelch ubiquitin ligase activity.

We were unable to conduct standard biochemical assays to determine whether HtsRC directly binds F-actin or nucleates actin polymerization. HtsRC is insoluble under native purification conditions and precipitates out of solution rapidly in the buffers used for *in vitro* F-actin biochemistry. Computational methods for predicting protein structure do not find any likely actin-binding motifs or other structural features in the HtsRC sequence, but instead indicate that this protein is mostly intrinsically disordered (Figure 1F) and that it likely becomes even more disordered following cleavage (not shown). It is still possible HtsRC contains an actin-binding motif; WH2 domains, one of the most ubiquitous actin binding domains, are found in regions of disorder (Carlier *et al*. 2011). Spire, a protein essential for oogenesis in *D. melanogaster*, is the founding member of a new class of actin nucleation promoting factors and contains four WH2 motifs in a region of disorder (Quinlan *et al*. 2005; Bradley *et al*. 2019). Although it has not been proven to be the case generally, there is evidence that intrinsically disordered proteins can become structured through interaction with a ligand, as is the case for Myelin Basic Protein (MBP) (Majava *et al*. 2010) and the G-actin binding protein Tarp (Tolchard *et al*. 2018). One avenue for future research is to determine if HtsRC becomes ordered in the presence of Filamin. We suspect that HtsRC’s intrinsic disorder and its propensity to form aggregates both *in vitro* and *in vivo* are functionally relevant.

### 3.2 Ring Canal Expansion and Stability

This study adds to a body of work in recent years reflecting the importance of cytoskeletal machinery in the expansion of the ring canals and in maintaining ring canal stability. Here, we have shown that HtsRC is a positive regulator of ring canal expansion. Overexpression promotes increased ring canal diameter, while lack of HtsRC results in decreased ring canal diameter. Furthermore, we see an increase in ring canal destabilization when HtsRC is absent.

Ring canal expansion starts once cysts have migrated to region 2b of the germarium and relies on the proper release of the constriction which occurred during cleavage furrow ingression (Ong and Tan 2010; Hamada-Kawaguchi *et al*. 2015). Although incomplete cytokinesis and ring canal expansion happen sequentially and Myosin activity is restricted to the mitotic phase of cyst formation, failure to regulate Myosin II during incomplete cytokinesis in *DMYPT* mutants results in ring canals that constrict too far and never expand at normally (Ong *et al*. 2010). Although the transition from cleavage furrow to ring canal is poorly understood, it may depend on proper regulation of yet unknown phospho-tyrosine species. Flies mutant for either *src64* or *tec29*, non-receptor tyrosine kinases known to localize to ring canals starting in the germarium, have defects in ring canal expansion despite normal recruitment and localization of Filamin, HtsRC, and Kelch (Guarnieri *et al*. 1998; Roulier *et al*. 1998; Dodson *et al*. 1998). In addition to showing a 30% decrease in ring canal diameter, which is comparable to the impact of both *htsRC* and Arp2/3 complex mutations (Hudson and Cooley 2002), *src64* mutants also exhibited frequent ring canal collapse (Dodson *et al*. 1998; O’Reilly *et al*. 2006). As we have described here for ring canal collapse in *htsRC*-specific mutants, collapse in *src64* mutants occured after an initial phase during which a subset of ring canals underwent some expansion and had an open lumen, but ultimately became unstable and collapsed into foci with no discernable lumen. Akap200, an anchoring protein for PKA regulatory subunits found at the outer rim of ring canals, has also been implicated in ring canal expansion but functions inversely to HtsRC and Src64 (Jackson and Berg 2002). In the absence of Akap200, ring canal expansion was increased and resulted in a thinner HtsRC inner rim, while overexpression of Akap200 resulted in smaller, thicker ring canals and both conditions caused ring canal collapse. Intriguingly, Akap200 and Src64 may function in the same pathway, as a single mutant Akap200 allele rescues ring canal size defects *src64* mutants. More recent work has revealed several other proteins that follow the same pattern as Akap 200. In particular, the knockdown of the formin Diaphanous or the Ste20 family kinase Misshapen promoted increased ring canal diameter and frequent ring canal collapse (Kline *et al*. 2018; Thestrup *et al*. 2020). Misshapen overexpression produces the inverse, decreased ring canal diameter.

The collapse of ring canals following both decreased and increased expansion suggests an exquisite balance between expansion and stability must be maintained in wildtype flies. When this balance is lost, ring canal stability is compromised. But how is the balance maintained in ring canals? HtsRC levels appear to correlate with the rate of ring canal expansion: decreased HtsRC levels lead to smaller ring canals, increased HtsRC levels lead to larger ring canals. In this work, we have shown that HtsRC is likely regulating ring canal expansion by regulating F-actin, but the mechanism by which the proteins listed above regulate ring canal expansion is less clear. The only known regulator of HtsRC, CRL3Kelch, regulates HtsRC protein but not ring canal expansion. One possible mechanism is that kinases like PKA (regulated by Akap200), Misshapen, Src64 and Tec29 impact ring canal expansion by directly or indirectly regulating the phosphorylation state and subsequent activity of HtsRC. HtsRC contains one or more phosphorylated residues (L. Petrella and L. Cooley, unpublished results) that may be important for regulating HtsRC activity at ring canals. Alternatively, the activity of these kinases may target other machinery for regulating F-actin once it has been recruited to the ring canal.

### 3.1 HtsRC evolution and fecundity

HtsRC is not essential for fertility, unlike other ring canal components including Kelch, Filamin and the Arp2/3 complex. What sets HtsRC apart from many other well studied components of female germline ring canals includes its status as a relatively young gene and its specificity to the female germline. To date, we have not found evidence of a HtsRC homolog outside of flies, although it is present in a recognizable form in the genomes of 26 species of *Drosophila*, as well as in house fly *(Musca domestica)* and in tsetse fly *(Glossina palpalis)* genomes (Figure S6). Comparison of HtsRC protein from these 28 species reveals regions of conservation interspersed with short, less conserved segments; the less conserved regions were often longer and more divergent in the two *non-Drosophila* species. Conserved sequence regions did not show any apparent correlation with regions of either order or disorder suggesting disordered regions are not less likely to be conserved (not shown). The exon encoding HtsRC many have been acquired in a fly lineage as recently as 60 million years ago, after the divergence of Dipteran flies. Although extensive work has not been done to confirm the sizes of ring canals in other insect species, we do know some insects have large ring canals while others do not (Haglund *et al*. 2011). Ovarian ring canals in bees can reach diameters of up to 4 microns (Ramamurty and Engels 1977; Patricio and Cruz-Landim 2006), while aphids appear have ring canals more similar to those in males (Pyka-Fosciak and Szklarzewicz 2008). Recent work observing ovaries in the butterfly *Pieris napi* suggests that they have actin rich ring canals in the female germline with diameters of around 10 microns (Mazurkiewicz-Kania *et al*. 2019). It is possible that differences in size could reflect the presence or absence of a HtsRC homolog.

Genes that are specific for reproduction exhibit different patterns of evolution compared to those in unrelated processes. Genes specific to reproduction are more likely to be rapidly evolving than the rest of the genome, especially those with testis-specific functions (Haerty *et al*. 2007). Although HtsRC is only expressed in the female germline, there is a testes-specific natural antisense transcript corresponding to the *htsRC* exon that may be blocking the expression of HtsRC in developing sperm. The *hts* locus additionally contains a poorly understood testis-specific splice variant, which also appears to be a recent evolutionary event in flies and is detectable in house flies and tsetse flies. We do not understand the function of either the non-coding RNA or the testis specific *hts* variant, but the presence of these transcripts hints at the complex evolutionary history of this gene and its regulation in both the male and female germline. Although the germline is permissive for the evolution of new genes or new gene functions, recent CRISPR-mediated mutagenesis of a subset of recently evolved genes determined that very few new genes are essential for fertility and none of those tested were essential for viability (Kondo *et al*. 2017).

Given that HtsRC is a newly evolved gene functioning in the germline, it is not surprising to find that it is not essential for either viability or fertility. Yet we have demonstrated an impact on fecundity--the presence of HtsRC confers an increase in the number of viable offspring a female can produce. This evidence supports the theory that the presence and expansion of a rigid F-actin cytoskeleton may enhance the functionality of ring canals to promote increased fecundity and may have been under positive selection. HtsRC is the product of a recent evolution event and is required to promote ring canal F-actin accumulation. Therefore, it is probable that the evolution of HtsRC in the fly lineage triggered changes in ring canal structure unique to the *Drosophila* female germline.

## Acknowledgements

We thank Nancy Pokrywka, David Ish-Horowicz, Mary Baylies, and Tian Xu for stocks used in this study. Stocks obtained from the Bloomington Drosophila Stock Center (NIH P40OD018537) were also used. We thank Mike Buszczak for sharing plasmids. We thank Tian Xu and Kaelyn Sumigray for providing access to the Leica SP8 confocal microscope. We thank Alexander Scherer and Anthony Koleske for their help with attempts to purify HtsRC for actin biochemistry. J.A.G. and K.M.M were funded in part by a Gruber Science Fellowship from Yale University and by NIH Training Grants [T32 GM007223 for K.M.M., T32 GM007499 for J.A.G.]. This work was supported by the NIH R01 GM043301 grant to LC.

## 4 Materials and Methods

### 4.1 Supplementary materials

Supplementary Figures PDF containing Figures S1-S6.

### 4.2 Reagent and Resource Sharing

Strains and plasmids are available upon request. The authors affirm that all data necessary for confirming the conclusions of the article are present within the article, figures, and tables. Further information and requests for resources and reagents should be directed to and will be fulfilled by the lead contact, Lynn Cooley.

### 4.3 Experimental Model

#### Drosophila genetics

Fruit flies were maintained at 22°C on standard fly food medium, unless otherwise indicated. Females were fattened for 8-24 hours on wet yeast past prior to dissection.

### 4.4 Method Details

#### Construction of *UASp-ovhts::GFP (K10UTR)* transgene

To make UASp-Ovhts::GFP, the coding sequence of Ovhts::GFP was PCR-amplified from UASH-Ovhts::GFP (contains *htsN4* 3’UTR; Petrella 2007) using primers to add Gateway recombination sequences to the 5’ and 3’ ends. PCR product encoding Ovhts::GFP was recombined into pDONR201, and a sequence-verified entry clone was recombined with pPW-attB, a UASp Gateway destination vector modified to include a phiC31 attB recombination site (gift from Mike Buszczak, UT Southwestern). The resulting pUASp-Ovhts::GFP plasmid was transformed into a strain containing the *attP2* landing site on chromosome III (68A4) at Rainbow Transgenics.

#### Construction of *UASH-ovhts::V5::APEX(htsN4UTR)* transgene

To make UASH-ovhts::V5::APEX, the coding sequence of APEX1 was PCR-amplified from pcDNA3-mito-APEX. The resulting PCR product contained a 5’ V5-tag linker sequence in addition to 5’XhoI and 3’ NotI cut sites. *pCOH-ovhts::GFP* (Petrella et al., 2007) was digested with XhoI and NotI to excise GFP and the V5::APEX1 fragment was ligated into the plasmid in-frame and in place of GFP. Transgenic flies were generated via P-element-mediated insertion at GenetiVision.

#### Analysis of MS/MS data for semi-tryptic peptides

Mass spectrometry data from Hudson *et al*. (2019) and Mannix *et al*. (2019) were searched for semi-tryptic peptides with up to two missed cleavages allowed. Briefly, mass spectrometry data were searched at MS Bioworks using Mascot (Matrix Science); searches were conducted against the UniProt *Drosophila melanogaster* database with common laboratory contaminants added as well as custom protein sequences for HtsRC that included known SNPs. The results were parsed into Scaffold (Proteome Software) for further analysis.

#### Ovary preparation and immunofluorescence

Ovaries were dissected in IMADS (Singleton and Woodruff 1994) or 1X PBS and fixed with 4% PFA (Electron Microscopy Sciences) in 1X PBS plus 0.3% Triton-X 100. For samples with actin cable imaging, we used a modified protocol based on Starble and Pokrywka (2018) with a 1 hour BSA incubation step. Primary antibody incubations were conducted for either 2 hours at room temperature or overnight at 4C and incubations with fluorescent secondaries plus TRITC-Phalloidin were done for 2 hours at room temperature. Antibody dilutions are indicated in the reagent table. Samples were mounted in ProLong Gold Antifade Mountant (Thermo Fisher Scientific) and allowed to cure 24-48 hours before imaging.

#### Microscopy

Samples were imaged on one of two microscope setups: a Leica SP8 scanning confocal system using a 40X Plan Apo 1.30 NA objective or 63X Plan APO 1.40 NA objective in combination with optical zoom (1.0x or 3.0x); or a Zeiss Axio Observer 7 inverted microscope with a CrEST X-Light V2 spinning disc system, Photometrics Prime BSi sCMOS camera, and a 40X C-Apo 1.2 NA water-immersion objective. Lower resolution egg images (Figure 4) were taken with a Leica MZFLIII stereo microscope (10X objective) fitted with a Leica MC120HD camera.

#### Image processing

All images shown or quantified were processed and analyzed with ImageJ/FIJI. All processing and treatments in ImageJ/FIJI were matched within each data set.

#### Egg chamber staging

Egg chamber stage was determined using both identifiable stage-specific features, such as shape, follicle cell behavior and oocyte size (Cummings and King 1969), and egg chamber area calculated for an ellipse based on the dimensions of the germ cells in a central z-slice, excluding the follicle cells (Jia *et al*. 2016). Where the two methods were in disagreement, visible features described in Cummings and King (1969) was given precedence.

#### Measurement of ring canal size

Ring canals were measured on z-projections using ImageJ/FIJI. The freehand line tool was used to plot the profile of a line bisecting the widest point of the ring canal, then the diameter was measured as the distance from the outside of one peak to the outside of the other at approximately 50% of the maximum fluorescence intensity for each peak. Ring canals were considered collapsed if no lumen was visible by eye or on a line scan of the ring canal, and diameter was measured at the half maximum for the single peak.

Violin plots were generated in Jupyter notebook using the matplotlib and seaborn visualization packages to display the distribution of ring canal diameter measurements across genotypes.

#### Fertility/fecundity tests

Fertility tests were performed in custom cages made from 14 mL polypropylene bacterial tubes attached in a honeycomb structure, which allowed tests to be performed on the same plates simultaneously for all genotypes. Cages were placed over grape juice or apple juice agar plates with charcoal added to increase visibility of eggs. One virgin and one male (always *w_1118_*) were mated in a single tube for 24 hours, then eggs were counted immediately (eggs laid) and 24 hours later (eggs remaining unhatched). Flies were initially left on plates for 24 hours to mature, then egg laying was measured for one to two and two to three-day old flies. Data presented is an aggregate of both days.

#### Measurement of mature eggs

Three virgins of each genotype and an equal number of wildtype *(w_1118_)* males were placed in mesh cages over grape juice agar plates with charcoal added to increase visibility of eggs. Plates were changed every 12 hours. Immediately after the 12-hour laying period, eggs on laying plates were imaged on a Leica MZFLIII stereo microscope (10X objective) fitted with a Leica MC120HD camera. Egg length was measured in ImageJ/FIJI.

#### Protein structure and disorder prediction

Prediction of intrinsically disordered regions within the Ovhts polyprotein were done using Spot-Disorder-Single (Hanson *et al*. 2018). The full length Ovhts polyprotein was used as the input. Disorder prediction was also run using the MetaDisorder (Kozlowski and Bujnicki 2012), which combines the results of multiple servers. Prediction of HtsRC structure and motifs was conducted using Phyre2 (Kelley *et al*. 2015).

#### Statistical tests

Statistical tests were conducted either in Prism 8 or in Microsoft Excel. Where appropriate, an F-test was conducted to determine if samples had equal variance, then the appropriate T-tests were used to determine statistical significance of the results. Individual tests used are indicated in figure legends. Where 3 samples were compared, an ANOVA was used and p-values displayed on graphs represent the results pairwise T-tests with p-values adjusted for multiple comparisons using built in statistical tests in Prism 8. Standard significance thresholds were used on graphs and are as follows: *, p < 0.05; **, p < 0.01; ***, p < 0.001; ****, p < 0.0001.

#### MARCM

MARCM stocks were generated by crossing *tubulin-Gal80 FRT40A* (2_nd_ chromosome) and *UAS-mCD8::GFP, tubulin-gal4* (3_rd_ chromosome) into a stock containing *hsFLP_1_* (X chromosome) to generate a balanced stock. Separately, an isogenized control *FRT40A* chromosome and the Arp2/3 complex alleles *(AprC1_Q29sd_ FRT40A* and *ArpC4_SH1036_ FRT40A)* on the second chromosome were crossed with *UASh-ovhts::V5::APEX (htsN4UTR)* on the 3_rd_ chromosome. The resulting stocks were crossed to the MARCM stock. Heat shocks were conducted on pupae for 1 hour at 38°C, then flies were allowed to develop for 6 days before dissection. Following heat shock, clones homozygous for Arp2/3 complex subunits or the control *FRT40A* chromosome were identified by the expression of CD8::GFP and the HtsRC::APEX transgene, both driven by *tubulin-Gal4*. The removal of any component of the Arp2/3 complex has previously been shown to inactivate the entire complex so mutant clones were considered null for Arp2/3 complex activity.

#### Quantification of HtsRC aggregates

Aggregates were characterized and quantified two ways: (1) using ImageJ/FIJI (Figure 6); (2) using Imaris 9.3 and Python code (Figure 7).

1. Z-projections of the follicle cell layer of stage 10B egg chambers were processed in ImageJ/FIJI. Positive F-actin foci were counted and measured in ImageJ/FIJI using the Auto-thresholding and Particle Analysis tools, with the minimum area of a foci set to 1 micron 2. Egg chamber area was calculated in FIJI using the Freehand and Area/Measure tools.
2. 3D image files were processed in Imaris 9.3.1 as follows: 150 pixel (24.375 micron) x 150 pixel regions of follicle cells were duplicated from images of stage 10A egg chambers using ImageJ; care was taken to choose regions with minimal curvature and three regions were chosen from approximately the same locations in each egg chamber. These images were imported into Imaris 9.3.1 and the Surfaces tool was used on the z-slices containing the somatic follicle cell layer. Aggregates (3D surfaces) were identified in the HtsRC (HtsRC antibody + Alexa 633) channel using background subtraction (local contrast) algorithm with automatic thresholding to distinguish objects (Surface Grain Size, 0.200 microns; Diameter of Largest Sphere, 0.200 microns). Nuclei were also identified by propagating the similar settings to the DAPI channel (Surface Grain Size, 0.325 microns; Diameter of Largest Sphere, 1.22 microns). Features of individual surfaces were exported including volume, center of mass, and sum of fluorescence intensity for all channels. Custom Python scripts in Jupyter notebook were used to perform calculations for each sample including density (summed fluorescence intensity/volume) and a ratio of F-actin intensity vs HtsRC. These calculations were always done on individual aggregates and averaged to generate a summary result for the 150×150-pixel region. A mean of three regions per egg chamber was used to generate plots shown in Figure 7; graphing was done using Prism 8. The relative Z-position of each aggregate was calculated as follows: the center of fluorescent intensity for DAPI signal was calculated for each 150×150 pixel region and used as the origin for that region and Z-positions closer than the origin to the coverslip were considered negative (basal side was always facing coverslip). Values for individual aggregates were sorted using a Python script, where individual aggregates were identified as apical, if they were positive relative to the nuclei, or basal, if they were negative relative to the nuclei.

#### Identification of HtsRC homologs in other species

Ovhts or just the HtsRC exon (*D. melanogaster)* were blasted against *Drosophila* as well as all available transcripts using NCBI blastn, blastx and blastp searches. *Musca domestica* sequence was identified by blasting *D. melanogaster* HtsRC against the other insect genomes using Vectorbase. Once identified, we repeated our search in Vectorbase using the *M. domestica* HtsRC sequence and identified the *G. palpalis* HtsRC homolog, which appears to be mis-annotated as an independent transcript but is likely part of an ovhts transcript. Once obtained, 28 HtsRC sequences were aligned using ClustalW and ClustalO algorithms in the Jalview software package. For this comparison, sequences matching the HtsRC exon were generated by removing all sequences corresponding to the HtsF/Adducin exons of *ovhts* and only these trimmed sequences were aligned.

## References

Andrianantoandro E., and T. D. Pollard, 2006 Mechanism of Actin Filament Turnover by Severing and Nucleation at Different Concentrations of ADF/Cofilin. Mol. Cell 24: 13–23. https://doi.org/10.1016/J.MOLCEL.2006.08.006

Bradley A. O., C. L. Vizcarra, H. M. Bailey, and M. E. Quinlan, 2019 Spire stimulates nucleation by Cappuccino and binds both ends of actin filaments. Mol. Biol. Cell 31: 273–286. https://doi.org/10.1091/mbc.E19-09-0550

Cant K., B. A. Knowles, M. S. Mooseker, and L. Cooley, 1994 Drosophila singed, a fascin homolog, is required for actin bundle formation during oogenesis and bristle extension. J. Cell Biol. 125: 369–380. https://doi.org/10.1083/jcb.125.2.369

Cant K., and L. Cooley, 1996 Single Amino Acid Mutations in Drosophila Fascin Disrupt Actin Bundling Function *in Vivo*; Genetics 143: 249.

Carlier M. F., V. Laurent, J. Santolini, R. Melki, D. Didry, et al., 1997 Actin depolymerizing factor (ADF/cofilin) enhances the rate of filament turnover: implication in actin-based motility. J. Cell Biol. 136: 1307–1322. https://doi.org/10.1083/jcb.136.6.1307

Carlier M.-F., C. Husson, L. Renault, and D. Didry, 2011 Control of Actin Assembly by the WH2 Domains and Their Multifunctional Tandem Repeats in Spire and Cordon-Bleu. Int. Rev. Cell Mol. Biol. 290: 55–85. https://doi.org/10.1016/B978-0-12-386037-8.00005-3

Cox R. T., and A. C. Spradling, 2003 A Balbiani body and the fusome mediate mitochondrial inheritance during*Drosophila*; oogenesis. Development 130: 1579. https://doi.org/10.1242/dev.00365

Cummings M. R., and R. C. King, 1969 The cytology of the vitellogenic stages of oogenesis in Drosophila melanogaster. I. General staging characteristics. J. Morphol. 128: 427–441. https://doi.org/10.1002/jmor.1051280404

Ding D., S. M. Parkhurst, and H. D. Lipshitz, 1993 Different genetic requirements for anterior RNA localization revealed by the distribution of Adducin-like transcripts during Drosophila oogenesis. Proc. Natl. Acad. Sci. U. S. A. 90: 2512–2516. https://doi.org/10.1073/pnas.90.6.2512

Dodson G. S., D. J. Guarnieri, and M. A. Simon, 1998 Src64 is required for ovarian ring canal morphogenesis during Drosophila oogenesis. Development 125: 2883.

Dominguez R., 2009 Actin filament nucleation and elongation factors--structure-function relationships. Crit. Rev. Biochem. Mol. Biol. 44: 351–366. https://doi.org/10.3109/10409230903277340

Greenbaum M. P., L. Ma, and M. M. Matzuk, 2007 Conversion of midbodies into germ cell intercellular bridges. Dev. Biol. 305: 389–396. https://doi.org/10.1016/j.ydbio.2007.02.025

Guarnieri D. J., G. S. Dodson, and M. A. Simon, 1998 SRC64 Regulates the Localization of a Tec-Family Kinase Required for Drosophila Ring Canal Growth. Mol. Cell 1: 831–840. https://doi.org/10.1016/S1097-2765(00)80082-9

Gutzeit H. O., 1990 The microfilament pattern in the somatic follicle cells of mid-vitellogenic ovarian follicles of Drosophila. Eur. J. Cell Biol. 53: 349–356.

Haerty W., S. Jagadeeshan, R. J. Kulathinal, A. Wong, K. Ravi Ram, et al., 2007 Evolution in the Fast Lane: Rapidly Evolving Sex-Related Genes in Drosophila. Genetics 177: 1321. https://doi.org/10.1534/genetics.107.078865

Haglund K., I. P. Nezis, and H. Stenmark, 2011 Structure and functions of stable intercellular bridges formed by incomplete cytokinesis during development. Commun. Integr. Biol. 4: 1–9.

Hamada-Kawaguchi N., Y. Nishida, and D. Yamamoto, 2015 Btk29A-mediated tyrosine phosphorylation of armadillo/β-catenin promotes ring canal growth in Drosophila oogenesis. PLoS One 10: e0121484–e0121484. https://doi.org/10.1371/journal.pone.0121484

Hanson J., K. Paliwal, and Y. Zhou, 2018 Accurate Single-Sequence Prediction of Protein Intrinsic Disorder by an Ensemble of Deep Recurrent and Convolutional Architectures. J. Chem. Inf. Model. 58: 2369–2376. https://doi.org/10.1021/acs.jcim.8b00636

Hinnant T. D., J. A. Merkle, and E. T. Ables, 2020 Coordinating Proliferation, Polarity, and Cell Fate in the Drosophila Female Germline. Front. Cell Dev. Biol. 8: 19.

Horne-Badovinac S., and D. Bilder, 2005 Mass transit: Epithelial morphogenesis in the Drosophila egg chamber. Dev. Dyn. 232: 559–574. https://doi.org/10.1002/dvdy.20286

Hudson A. M., and L. Cooley, 2002 A subset of dynamic actin rearrangements in Drosophila requires the Arp2/3 complex. J. Cell Biol. 156: 677–687. https://doi.org/10.1083/jcb.200109065

Hudson A. M., K. M. Mannix, J. A. Gerdes, M. C. Kottemann, and L. Cooley, 2019 Targeted substrate degradation by Kelch controls the actin cytoskeleton during ring canal expansion. Development 146: dev169219. https://doi.org/10.1242/dev.169219

Huelsmann S., J. Ylänne, and N. H. Brown, 2013 Filopodia-like Actin Cables Position Nuclei in Association with Perinuclear Actin in Drosophila Nurse Cells. Dev. Cell 26: 604–615. https://doi.org/10.1016/J.DEVCEL.2013.08.014

Jackson S. M., and C. A. Berg, 2002 An A-kinase anchoring protein is required for Protein kinase A regulatory subunit localization and morphology of actin structures during oogenesis in *Drosophila*; Development 129: 4423.

Jia D., Q. Xu, Q. Xie, W. Mio, and W.-M. Deng, 2016 Automatic stage identification of Drosophila egg chamber based on DAPI images. Sci. Rep. 6: 18850. https://doi.org/10.1038/srep18850

Kelley L. A., S. Mezulis, C. M. Yates, M. N. Wass, and M. J. E. Sternberg, 2015 The Phyre2 web portal for protein modeling, prediction and analysis. Nat. Protoc. 10: 845–858. https://doi.org/10.1038/nprot.2015.053

Kelso R. J., A. M. Hudson, and L. Cooley, 2002 Drosophila Kelch regulates actin organization via Src64-dependent tyrosine phosphorylation. J. Cell Biol. 156: 703–713. https://doi.org/10.1083/jcb.200110063

Kline A., T. Curry, and L. Lewellyn, 2018 The Misshapen kinase regulates the size and stability of the germline ring canals in the Drosophila egg chamber. Dev. Biol. 440: 99–112. https://doi.org/10.1016/J.YDBIO.2018.05.006

Kondo S., J. Vedanayagam, J. Mohammed, S. Eizadshenass, L. Kan, et al., 2017 New genes often acquire male-specific functions but rarely become essential in Drosophila. Genes Dev. 31: 1841–1846. https://doi.org/10.1101/gad.303131.117

Kozlowski L. P., and J. M. Bujnicki, 2012 MetaDisorder: a meta-server for the prediction of intrinsic disorder in proteins. BMC Bioinformatics 13: 111.

Lee T., and L. Luo, 1999 Mosaic Analysis with a Repressible Cell Marker for Studies of Gene Function in Neuronal Morphogenesis. Neuron 22: 451–461. https://doi.org/10.1016/S0896-6273(00)80701-1

Lin H., L. Yue, and A. C. Spradling, 1994 The Drosophila fusome, a germline-specific organelle, contains membrane skeletal proteins and functions in cyst formation. Development 120: 947.

Loyer N., I. Kolotuev, M. Pinot, and R. Le Borgne, 2015 Drosophila E-cadherin is required for the maintenance of ring canals anchoring to mechanically withstand tissue growth. Proc. Natl. Acad. Sci. 112: 12717. https://doi.org/10.1073/pnas.1504455112

Lu K., L. Jensen, L. Lei, and Y. M. Yamashita, 2017 Stay Connected: A Germ Cell Strategy. Trends Genet. 33: 971–978. https://doi.org/10.1016/j.tig.2017.09.001

Mahajan-Miklos S., and L. Cooley, 1994 The villin-like protein encoded by the Drosophila quail gene is required for actin bundle assembly during oogenesis. Cell 78: 291–301. https://doi.org/10.1016/0092-8674(94)90298-4

Majava V., C. Wang, M. Myllykoski, S. M. Kangas, S. U. Kang, et al., 2010 Structural analysis of the complex between calmodulin and full-length myelin basic protein, an intrinsically disordered molecule. Amino Acids 39: 59–71. https://doi.org/10.1007/s00726-009-0364-2

Mannix K. M., R. M. Starble, R. S. Kaufman, and L. Cooley, 2019 Proximity labeling reveals novel interactomes in live *Drosophila*; tissue. Development 146: dev176644. https://doi.org/10.1242/dev.176644

Mazurkiewicz-Kania M., B. Simiczyjew, and I. Jędrzejowska, 2019 Differentiation of follicular epithelium in polytrophic ovaries of Pieris napi (Lepidoptera: Pieridae) —how far to Drosophila model. Protoplasma 256: 1433–1447. https://doi.org/10.1007/s00709-019-01391-1

McLean P. F., and L. Cooley, 2013 Protein Equilibration Through Somatic Ring Canals in *Drosophila*; Science (80-.). 340: 1445. https://doi.org/10.1126/science.1234887

Mullins R. D., J. A. Heuser, and T. D. Pollard, 1998 The interaction of Arp2/3 complex with actin: nucleation, high affinity pointed end capping, and formation of branching networks of filaments. Proc. Natl. Acad. Sci. U. S. A. 95: 6181–6186. https://doi.org/10.1073/pnas.95.11.6181

Nakamura F., T. M. Osborn, C. A. Hartemink, J. H. Hartwig, and T. P. Stossel, 2007 Structural basis of filamin A functions. J. Cell Biol. 179: 1011–1025. https://doi.org/10.1083/jcb.200707073

Nakamura F., T. P. Stossel, and J. H. Hartwig, 2011 The filamins: organizers of cell structure and function. Cell Adh. Migr. 5: 160–169. https://doi.org/10.4161/cam.5.2.14401

Ngo K. X., N. Kodera, E. Katayama, T. Ando, and T. Q. P. Uyeda, 2015 Cofilin-induced unidirectional cooperative conformational changes in actin filaments revealed by highspeed atomic force microscopy. Elife 4: e04806. https://doi.org/10.7554/eLife.04806

O’Reilly A. M., A. C. Ballew, B. Miyazawa, H. Stocker, E. Hafen, et al., 2006 Csk differentially regulates Src64 during distinct morphological events in Drosophila germ cells. Development 133: 2627–2638.

Ong S., and C. Tan, 2010 Germline cyst formation and incomplete cytokinesis during Drosophila melanogaster oogenesis. Dev. Biol. 337: 84–98. https://doi.org/10.1016/J.YDBIO.2009.10.018

Ong S., C. Foote, and C. Tan, 2010 Mutations of DMYPT cause over constriction of contractile rings and ring canals during Drosophila germline cyst formation. Dev. Biol. 346: 161–169. https://doi.org/10.1016/J.YDBIO.2010.06.008

Patricio K., and C. Cruz-Landim, 2006 Ultrastructural aspects of the intercellular bridges between female bee germ cells. Braz. J. Biol. 66: 309–315.

Petrella L. N., T. Smith-Leiker, and L. Cooley, 2007 The Ovhts polyprotein is cleaved to produce fusome and ring canal proteins required for Drosophila oogenesis. Development 134: 703–712. https://doi.org/10.1242/dev.02766

Pokrywka N. J., H. Zhang, and K. Raley-Susman, 2014 Distinct roles for hu li tai shao and swallow in cytoskeletal organization during Drosophila oogenesis. Dev. Dyn. 243: 906–916. https://doi.org/10.1002/dvdy.24132

Pyka-Fosciak G., and T. Szklarzewicz, 2008 Germ cell cluster formation and ovariole structure in viviparous and oviparous generations of the aphid Stomaphis quercus. Int. J. Dev. Biol. 52: 259–265.

Quinlan M. E., J. E. Heuser, E. Kerkhoff, and R. Dyche Mullins, 2005 Drosophila Spire is an actin nucleation factor. Nature 433: 382–388. https://doi.org/10.1038/nature03241

Ramamurty P. S., and W. Engels, 1977 Occurrence of intercellular bridges between follicle epithelial cells in the ovary of Apis mellifica queens. J. Cell Sci. 24: 195.

Razinia Z., T. Mäkelä, J. Ylänne, and D. A. Calderwood, 2012 Filamins in Mechanosensing and Signaling. Annu. Rev. Biophys. 41: 227–246. https://doi.org/10.1146/annurev-biophys-050511-102252

Robinson D. N., K. Cant, and L. Cooley, 1994 Morphogenesis of Drosophila ovarian ring canals. Development 120: 2015.

Robinson D. N., and L. Cooley, 1996 Stable intercellular bridges in development: the cytoskeleton lining the tunnel. Trends Cell Biol. 6: 474–479. https://doi.org/10.1016/0962-8924(96)84945-2

Robinson D. N., T. A. Smith-Leiker, N. S. Sokol, A. M. Hudson, and L. Cooley, 1997 Formation of the Drosophila ovarian ring canal inner rim depends on cheerio. Genetics 145: 1063–1072.

Roulier E. M., S. Panzer, and S. K. Beckendorf, 1998 The Tec29 Tyrosine Kinase Is Required during Drosophila Embryogenesis and Interacts with Src64 in Ring Canal Development. Mol. Cell 1: 819–829. https://doi.org/10.1016/S1097-2765(00)80081-7

Singleton K., and R. I. Woodruff, 1994 The Osmolarity of Adult Drosophila Hemolymph and Its Effect on Oocyte-Nurse Cell Electrical Polarity. Dev. Biol. 161: 154–167. https://doi.org/10.1006/DBIO.1994.1017

Sokol N. S., and L. Cooley, 1999 Drosophila Filamin encoded by the cheerio locus is a component of ovarian ring canals. Curr. Biol. 9: 1221–1230. https://doi.org/10.1016/S0960-9822(99)80502-8

Spracklen A. J., D. J. Kelpsch, X. Chen, C. N. Spracklen, and T. L. Tootle, 2014 Prostaglandins temporally regulate cytoplasmic actin bundle formation during Drosophila oogenesis. Mol. Biol. Cell 25: 397–411. https://doi.org/10.1091/mbc.E13-07-0366

Starble R., and N. J. Pokrywka, 2018 The retromer subunit Vps26 mediates Notch signaling during Drosophila oogenesis. Mech. Dev. 149: 1–8. https://doi.org/10.1016/J.MOD.2017.10.001

Thestrup J., M. Tipold, A. Kindred, K. Stark, T. Curry, et al., 2020 The Arp2/3 complex and the formin, Diaphanous, are both required to regulate the size of germline ring canals in the developing egg chamber. Dev. Biol. https://doi.org/10.1016/J.YDBIO.2020.01.007

Tilney L. G., M. S. Tilney, and G. M. Guild, 1996 Formation of actin filament bundles in the ring canals of developing Drosophila follicles. J. Cell Biol. 133: 61–74. https://doi.org/10.1083/jcb.133.1.61

Tolchard J., S. J. Walpole, A. J. Miles, R. Maytum, L. A. Eaglen, et al., 2018 The intrinsically disordered Tarp protein from chlamydia binds actin with a partially preformed helix. Sci. Rep. 8: 1960. https://doi.org/10.1038/s41598-018-20290-8

Whittaker K. L., D. Ding, W. W. Fisher, and H. D. Lipshitz, 1999 Different 3’ untranslated regions target alternatively processed hu-li tai shao (hts) transcripts to distinct cytoplasmic locations during Drosophila oogenesis. J. Cell Sci. 112: 3385.

Wilson P. G., 1999 BimC motor protein KLP61F cycles between mitotic spindles and fusomes in Drosophila germ cells. Curr. Biol. 9: 923–S3. https://doi.org/10.1016/S0960-9822(99)80400-X

Xue F., and L. Cooley, 1993 Kelch encodes a component of intercellular bridges in Drosophila egg chambers. Cell 72: 681–693. https://doi.org/10.1016/0092-8674(93)90397-9

Yue L., and A. C. Spradling, 1992 hu-li tai shao, a gene required for ring canal formation during Drosophila oogenesis, encodes a homolog of adducin. Genes Dev. 6: 2443–2454. https://doi.org/10.1101/gad.6.12b.2443

Zallen J. A., Y. Cohen, A. M. Hudson, L. Cooley, E. Wieschaus, et al., 2002 SCAR is a primary regulator of Arp2/3-dependent morphological events in Drosophila. J. Cell Biol. 156: 689–701. https://doi.org/10.1083/jcb.200109057

