## Supplemental figures S1-S6 for "HtsRC-mediated accumulation of F-actin regulates ring canal size during Drosophila melanogaster oogenesis"

**Supplementary figures S1-S6 for:**

### HtsRC-mediated accumulation of F-actin regulates ring canal size during Drosophila melanogaster oogenesis

Julianne A. Gerdes*, Katelynn M. Mannix*, Andrew M. Hudson*, Lynn Cooley*,†,‡,§

**Figure S1: Ring canals connecting nurse cells to oocytes retain more F-actin than nurse cell ring canals.** Representative ring canals from several stages labeled with Phalloidin stain (magenta) and Filamin antibody (green). (left) The ring canals connecting pairs of nurse cells never form a robust F-actin cytoskeleton although Filamin localizes normally. (right) The ring canals connecting nurse cells to the oocyte form an actin cytoskeleton at later stages (9 and 10) which partially colocalizes with the Filamin stain marking the inner rim. Arrows point to the inner rim to allow comparison of Filamin and F-actin channels.

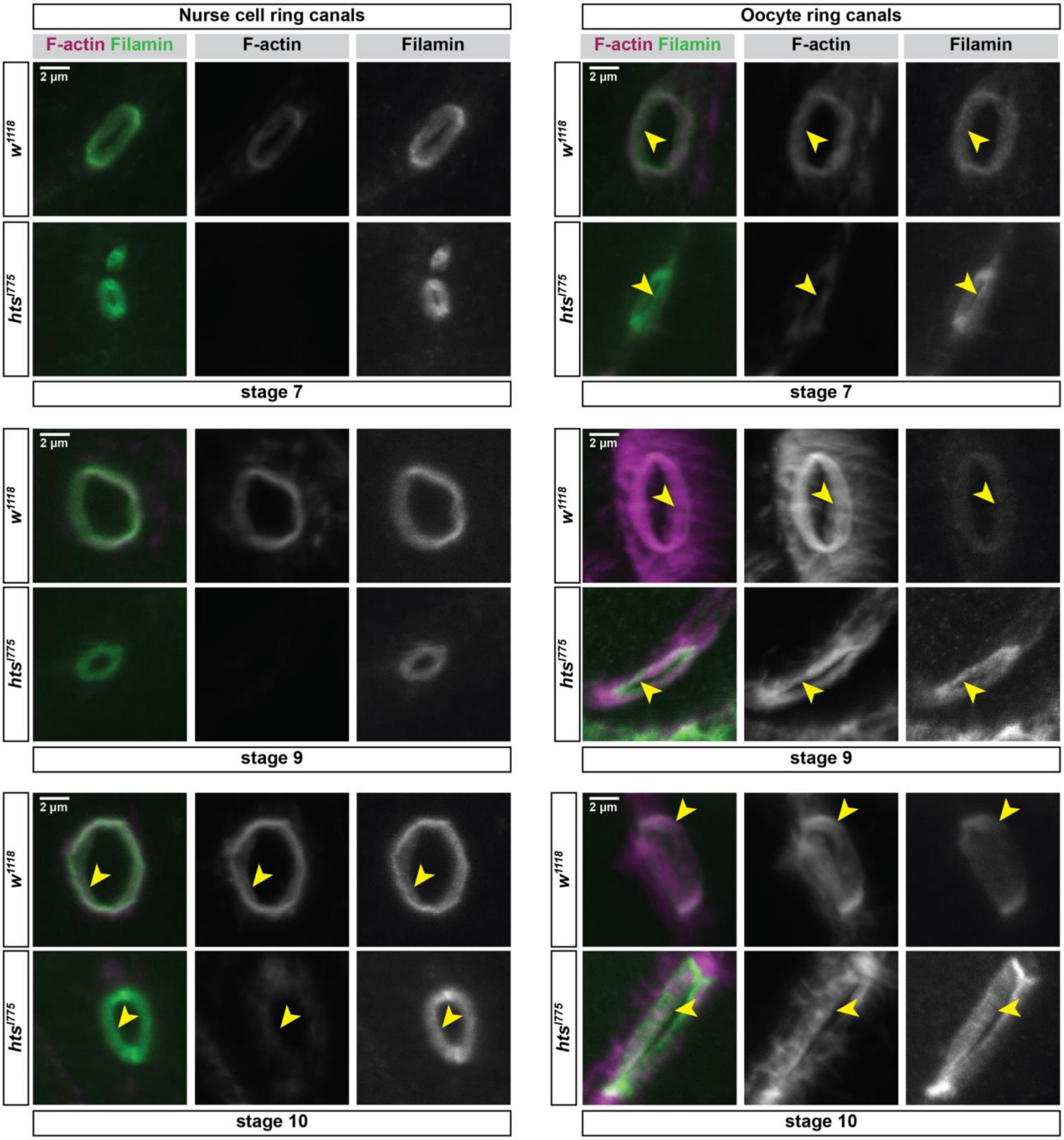

**Figure S2: HtsRC persists on actin cables in *cheerio* (Filamin) mutants**

(A) Diagram of the *cheerio* (Filamin) gene and its protein products. Two independent transcription start sites promote the transcription of a full length Filamin (Filamin240) and a short Filamin (Filamin90). *cher1* is an allele with no known lesion that does not express detectable Filamin protein in the germline or somatic follicle cells. *cherM113* and *cherM919* were a gift from the David Ish-Horowicz lab. *cherM113* causes a nonsense mutation in the actin binding domain, while *cherM919* contains a point mutation which converts an AG to TG and eliminates a splice acceptor. *cherMI07480-GFSTF.0* is an intronic insertion allele from the Mimic collection which has been swapped for a GFP containing cassette. GFP protein is inserted in the hinge of Filamin240 and roughly 70% or Filamin protein in ovary lysates contains GFP (results not shown). (B) HtsRC (green) and F-actin (magenta) stain of *w1118* control compared to three *cheerio* mutants. HtsRC localizes to actin cables and ring canals in both wild type and to cables in *cher* mutants. Antibody to the C-terminus of Filamin (not included in the merge) labels ring canals but does not stain actin cables.

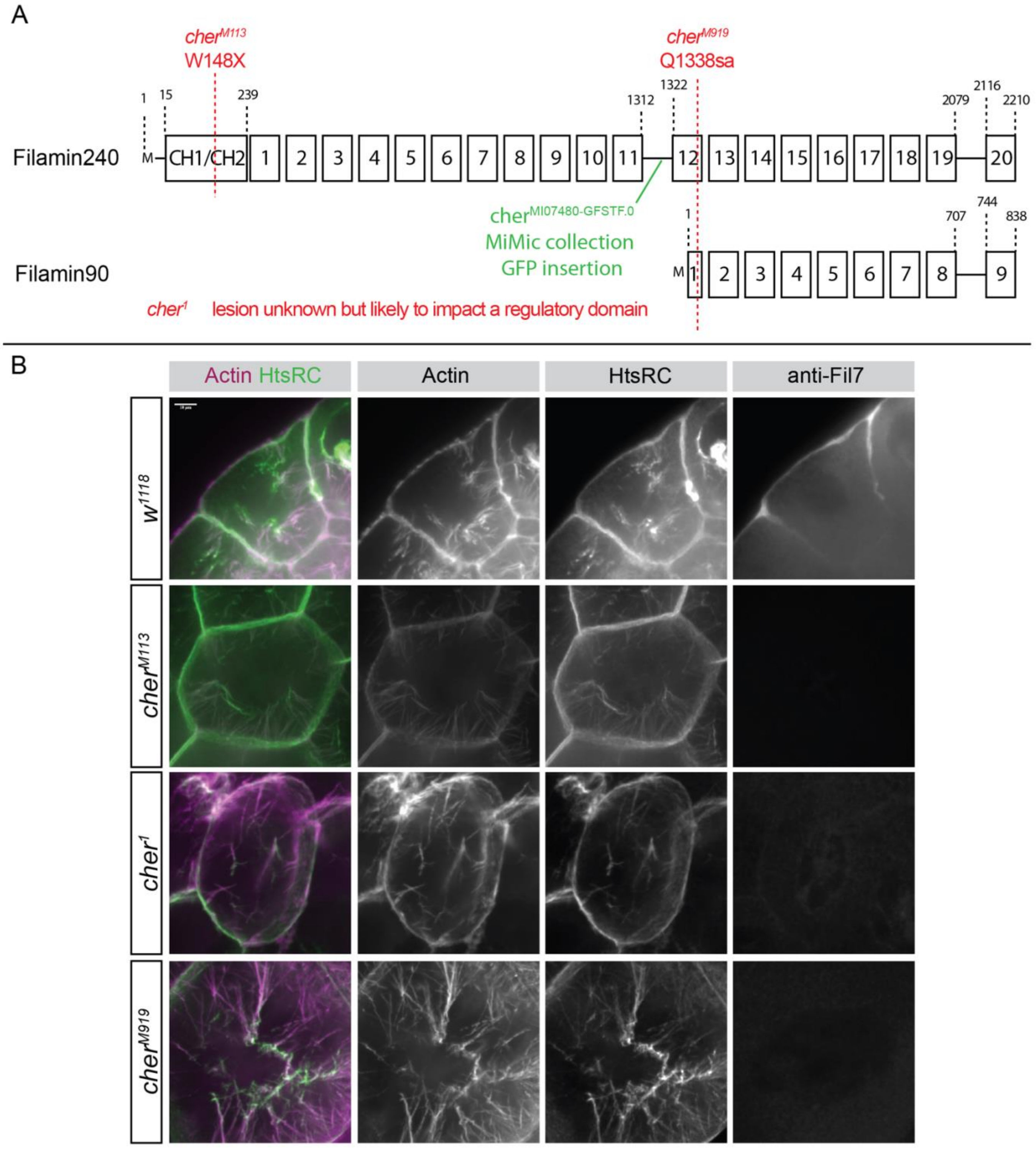

**Figure S3: HtsRC/F-actin aggregates formation is consistent across various transgenes and can be regulated by Kelch.** (A-E) View of somatic follicle cells from the basal end in *w1118* control (A) compared with four different *ovhts* transgenes. (F-J) View of somatic follicle cells from the side with the basal end oriented downward and the apical end/germline oriented upward. (K-M) *UASh-ovhts::GFP-htsN4* transgene (same as C, H) driven in somatic follicle cells under the control of the hindsight-Gal4 driver. An mCherry control (K), mCherry::Kelch (L) or unregulated Kelch (M) is coexpressed with the same driver. (N-Q) Quantification of aggregate/foci number. Individual data points represent measurements from individual egg chambers. Significance was determined using one-way ANOVA, Tukey's multiple comparison test. significance thresholds as follows: p<0.05(*); p<0.0001(****). (N), aggregate size (O), total summed aggregate area (P) and total summed aggregate fluorescence for all three genotypes.

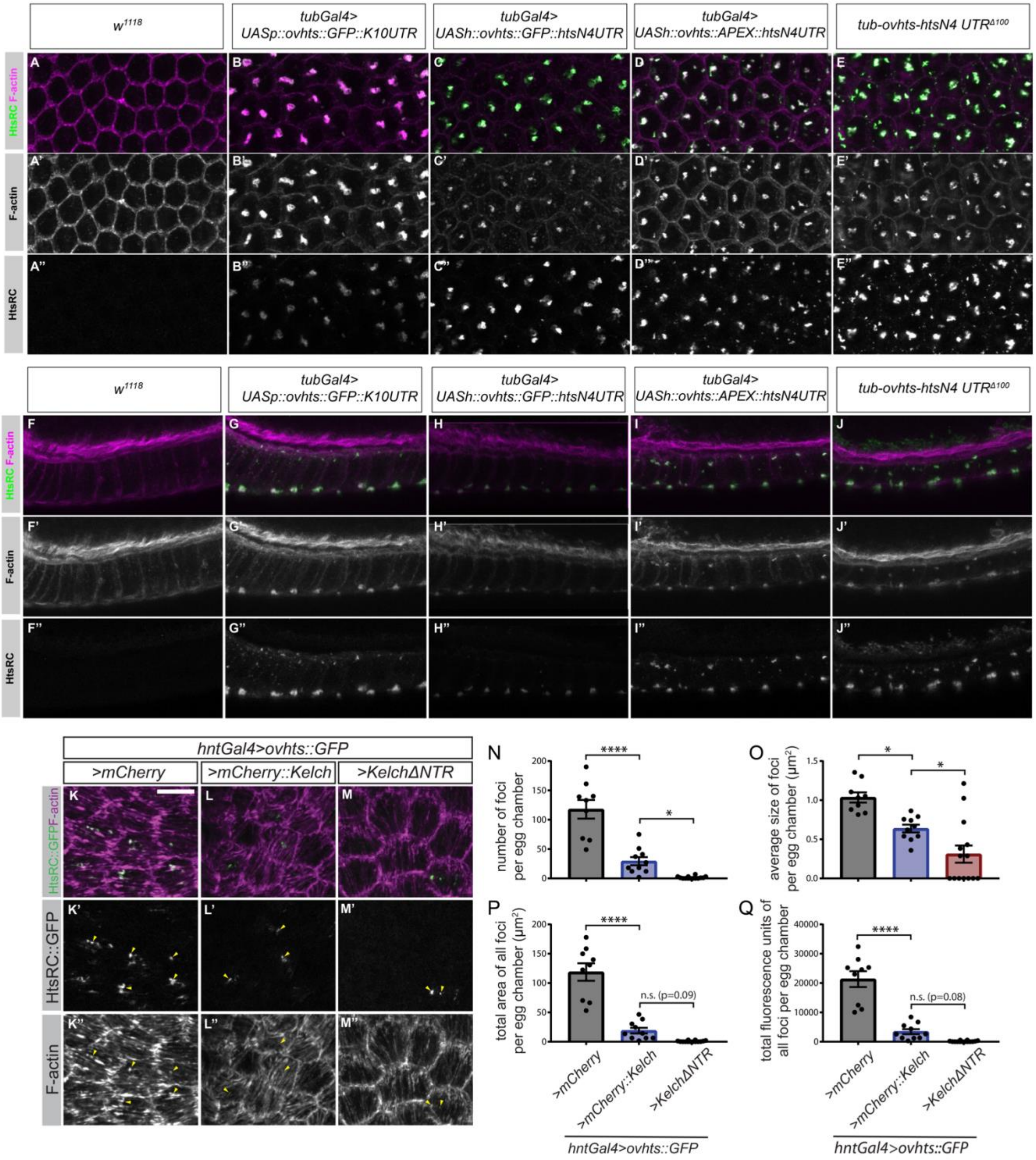

Figure S4: **Ectopic HtsRC expression drives F-actin actin aggregate formation independent of Arp2/3 complex activity.** Heatshocks were performed on flies heterozygous for *FRT40A Tubulin-Gal80* and either a wildtype control FRT40A chromosome or an FRT40A chromosome containing a mutant allele. Clones expressing two copies of the control allele or two copies of the mutant allele were marked by the expression of CD8::GFP (blue) and HtsRC::V5::APEX (green) driven by *tubulin-Gal4* expression. Neighboring cells (no GFP) contained either one or two copies of *tubulin-Gal80* and did not express GFP or HtsRC protein. Samples were also stained with Phalloidin. (A) Control clone (B) ArpC1$_{Q29sd}$ mutant clone (C) ArpC4$_{SH1036}$ mutant clone.

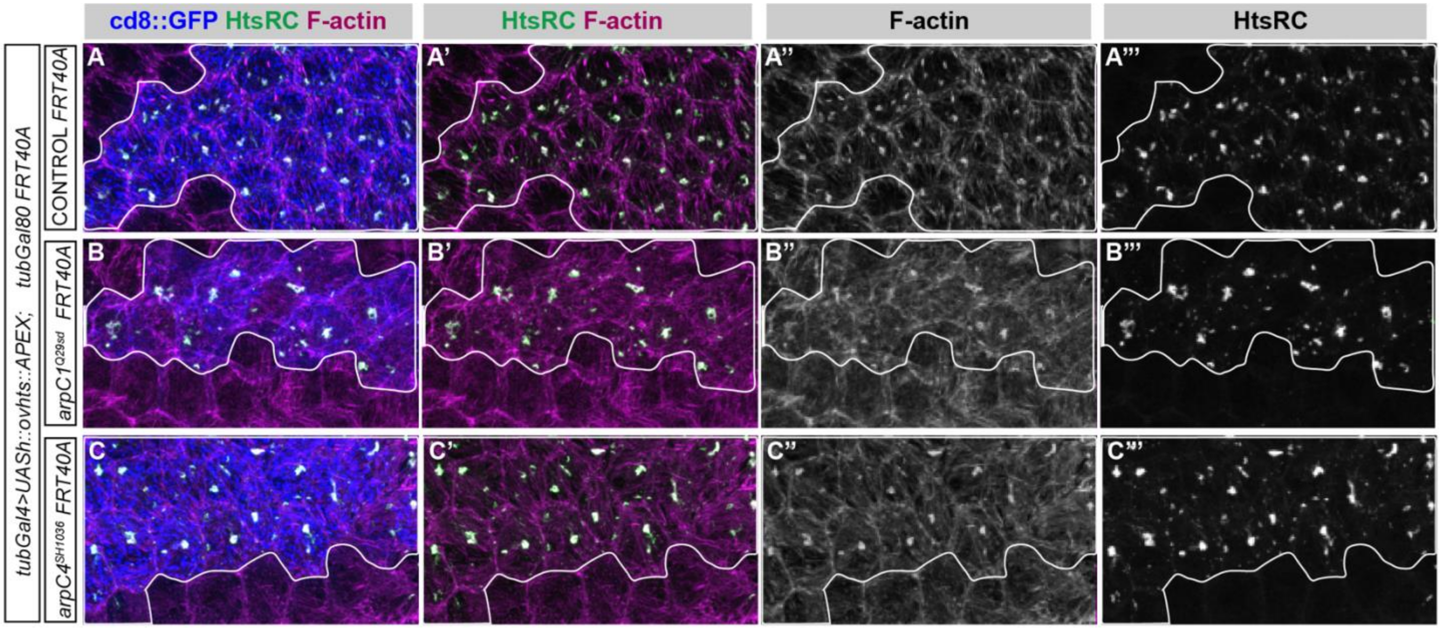

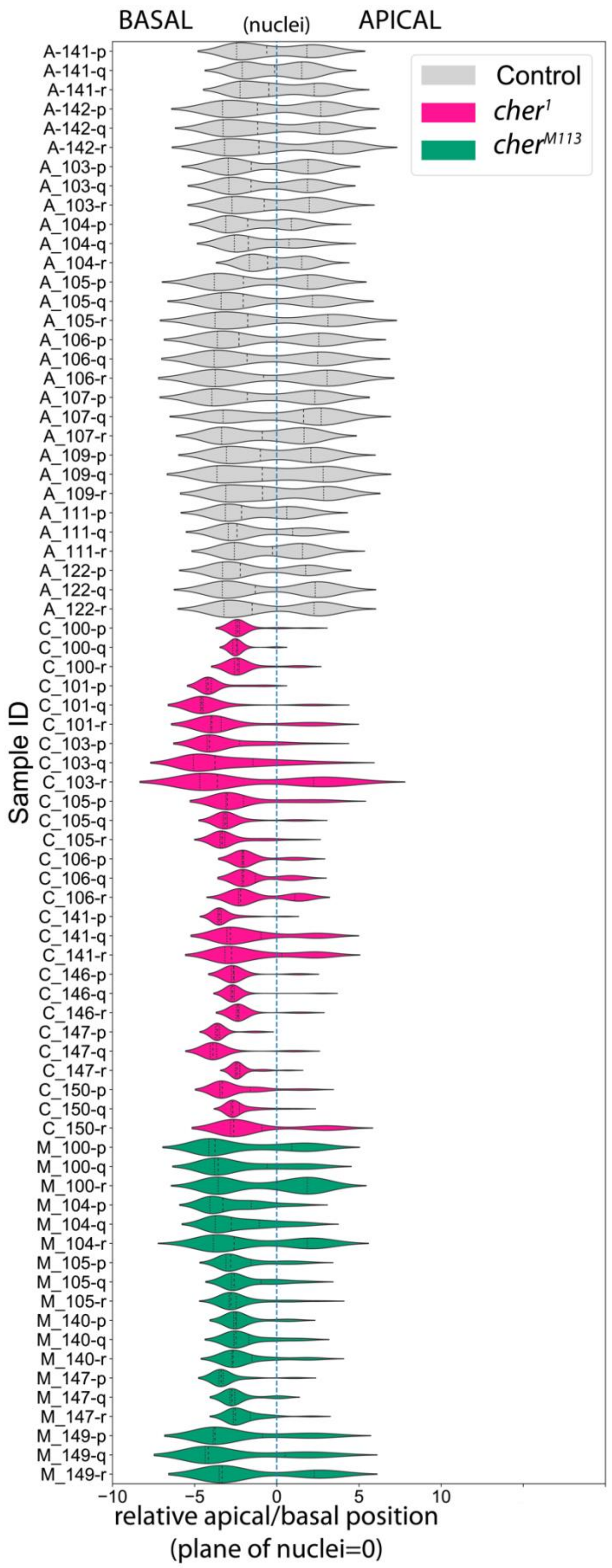


**Figure S5: HtsRC/F-actin aggregates rarely form at the apical surface in *cheerio* mutants.** Aggregate location within follicle cells was determined in Imaris for all aggregates within three regions of interest (regions p, q, and r; 24x24 micron squares) per egg chamber. Z-position at the center of mass (based on HtsRC intensity) for each aggregate was used as a proxy for the location of the entire aggregate or cluster of aggregates; aggregates in close proximity were reported by Imaris as a single aggregate. The position of the nuclei was calculated using the center of mass for DAPI stain and used to normalize the relative aggregate position between samples. Although the x and y dimensions of the regions of interest were fixed, the z dimension varied with the depth of the follicle cell layer.

**Supplemental Figure 6 (on following page): Predicted HtsRC homologs are present in 28 fly species.** Clustal Omega alignment of HtsRC proteins from 26 *Drosophila* species as well as *Musca domestica* and *Glossina palpalis.* Gaps are indicated by blank space. Residue conservation within this group is indicated by black boxes under the alignment. Conserved residues are colored according to the Clustal X color scheme with non conserved residues left white: hydrophobic (blue), positive charge (red), negative charge (magenta), polar (green),cysteines (pink), glycines (orange), prolines (yellow), and aromatics (cyan). The site of Ovhts polyprotein cleavage and Kelch binding found in *D. melanogaster* are indicated by black outlines. *D. melanogaster* sequence is indicated by a red dashed line. Species in alignment (in order) are as follows: *D. arizonae, D. biarmipes, D. bipectinata, D. busckii, D. elegans, D. erecta, D. eugracilis, D. ficusphila, D. grimshawi, D. hydei, D. kikkawai, D. melanogaster, D. miranda, D. mojavensis, D. navojoa, D. persimilis, D. pseudoobscura, D. rhopaloa, D. sechellia, D. serrata, D. simulans, D. suzukii, D. takahasii, D. virilis, D. willistoni, D. yakuba, G. palpalis, M. domestica.*

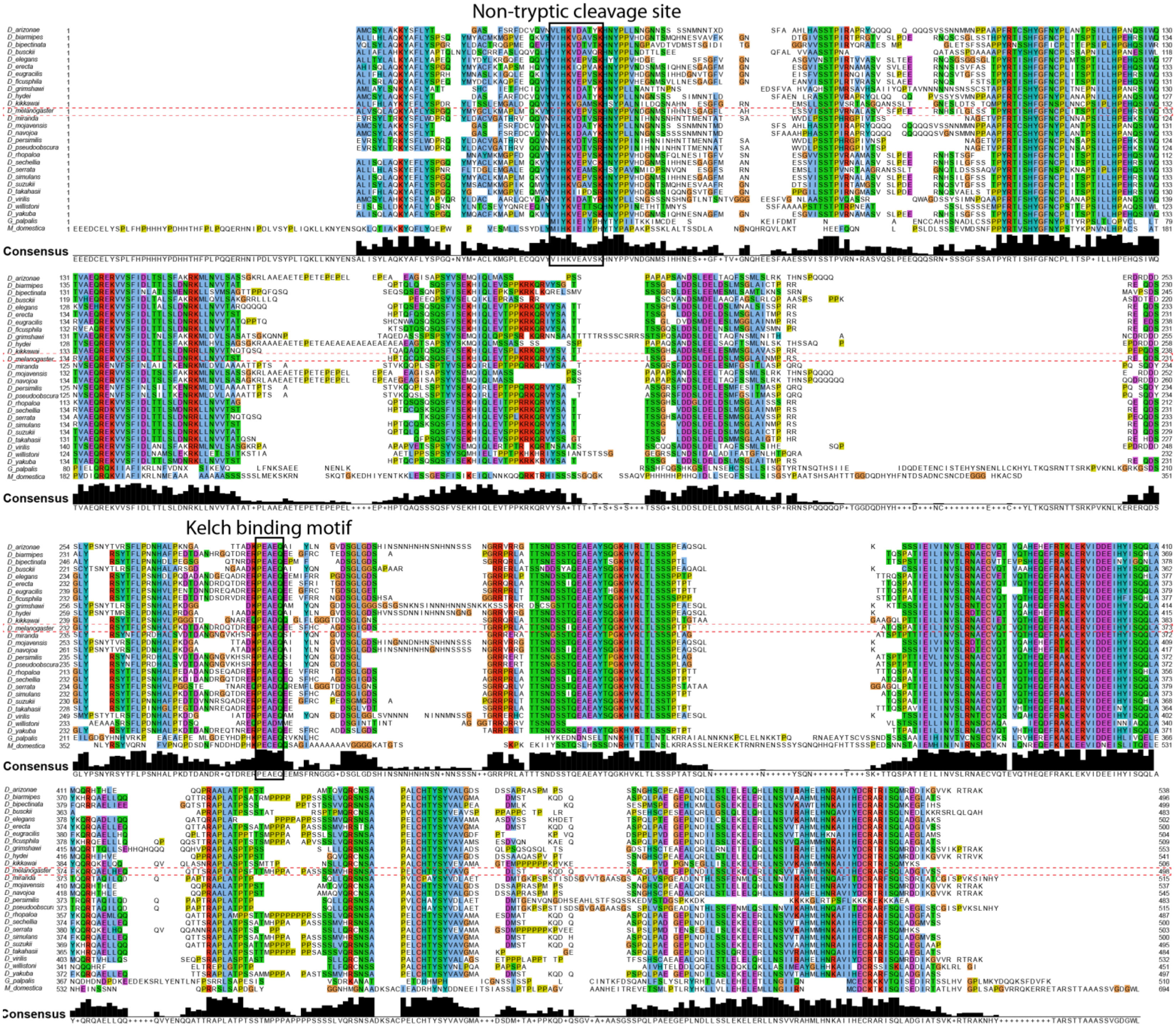
Non-tryptic cleavage site
Kelch binding motif
Consensus
